# Viral genetic variation programs divergent neuroimmune states during rabies infection

**DOI:** 10.64898/2026.11.20.768130

**Authors:** Sebastian Gaßmann, Rachel Mullner, Luca M. Zaeck, Conrad M. Freuling, Laura Schmid, Laura Rzepa, Ola Bagato, Thomas Müller, Falk Butter, Stefan Finke, Dmitry S. Ushakov

## Abstract

Viral burden does not necessarily predict the extent or composition of the host immune response, reflecting the complexity of virus - host interactions. Rabies virus (RABV) provides a powerful model to investigate this question because genetically distinct RABVs differ in replication, immune evasion, and neuroimmune activation. Here, we performed integrated temporal proteomic and spatial imaging analyses of recombinant RABV, including pathogenic, attenuated, and a newly generated interferon-sensitive variant with disrupted phosphoprotein antagonistic activity. Following intracerebral infection of mice, viruses established distinct neuroimmune states that diverged over time in antiviral signaling, complement activation, and immune response. Notably, the pathogenic field viruses rDogA and rFox both exhibited substantial viral accumulation, but elicited markedly different central nervous system (CNS) responses, with rDogA inducing limited host immune engagement, whereas rFox triggered early and broad antiviral, complement, and immune recruitment programs. Disruption of phosphoprotein-mediated interferon antagonism in rDogA enhanced interferon-stimulated gene expression and antiviral effectors, restricting viral accumulation without inducing broad neuroimmune remodeling. Conversely, the attenuated glycoprotein-modified rSAD GASGAS elicited strong complement activation and T cell recruitment with little detectable viral levels, revealing a highly immunogenic neuroimmune state. Spatial profiling further identified virus-dependent patterns of myeloid, astrocyte-associated, complement, and lymphocyte responses across brain regions. Together, these findings demonstrate how viral genetic background and its modulation of host antiviral responses shape divergent neuroimmune states in which viral accumulation, antiviral immunity, and CNS immune activation are differentially coordinated.

## Introduction

Neurotropic viruses face a unique challenge in balancing their replication with avoidance of immune control within the central nervous system (CNS) (1, 2). Although insufficient antiviral responses can permit viral replication and dissemination, excessive immune activation may contribute to neurological pathology (3). This balance is particularly evident in CNS infections caused by viruses such as rabies virus (RABV), West Nile virus, and herpesviruses, where viral and host determinants shape the magnitude and timing of antiviral responses. For neurotropic flaviviruses such as West Nile virus, innate immune responses are essential for restricting infection but can also contribute to neurological damage when dysregulated (4). Conversely, impaired innate antiviral sensing can result in uncontrolled CNS infection, as demonstrated by susceptibility to Herpes simplex virus type 1 encephalitis in individuals with defects in TLR3-dependent interferon response (5). However, how viral genetic variation programs distinct immune trajectories within infected neural tissue remains poorly understood.

RABV exhibits substantial variation in its interactions with host cells and immune responses across field isolates, cell culture-adapted viruses, and attenuated vaccine viruses, despite a highly conserved genome organization, providing a powerful model for dissecting the genetic and immunological underpinnings of neurovirulence (6, 7). Field isolates, such as dog- and fox-derived RABV circulating in natural reservoirs, are characterized by efficient neuroinvasion, limited cytopathology, and a remarkable capacity to evade early host immune detection (8). These viruses can replicate in neurons while delaying or modulating early host immune detection, allowing the CNS infection to progress before effective adaptive immune responses are established (9–11).

In contrast, laboratory-adapted and attenuated RABV strains contain defined genetic alterations that influence viral replication dynamics, neurovirulence, and host immune engagement (8, 12). Laboratory strains such as the challenge virus standard (CVS-11) retain many pathogenic features and serve as experimental models of virulent infection, whereas attenuated Street Alabama Dufferin (SAD) lineage viruses represent vaccine-adapted variants in which genetic changes modify virus - host interactions and infection outcome (12, 13).

At the molecular level, viral glycoprotein (G) is a central determinant of neurovirulence and immune engagement (14, 15). Field viruses, CVS-11, and early attenuated SAD derivatives such as SAD L16 (16) retain a conserved basic residue at position 333 of G that is essential for efficient neuroinvasion (17, 18). Importantly, the SAD L16 is attenuated following intramuscular administration, but retains high neurovirulence after direct CNS inoculation in mice, indicating that attenuation is primarily associated with restricted neuroinvasion and host-specific determinants rather than an inability to replicate in the CNS (19, 20). Later SAD-derived vaccine strains, such as SAG2 and SPBN GASGAS variants carrying glycoprotein modifications including either single G333E (SAG2), or double G333E and N194S substitutions (SPBN GASGAS), display reduced neurovirulence and enhanced immunogenicity (15, 21, 22). However, additional viral proteins also contribute to virus-specific phenotypes. The phosphoprotein (P), a key antagonist of type I interferon signaling, supports suppression of innate antiviral responses (23, 24). Although less variable, nucleoprotein (N), matrix protein (M), and viral polymerase (L) contribute to replication efficiency, virion assembly, and viral fitness, potentially influencing antigen availability and downstream immune activation (9, 25, 26). Moreover, the N protein acts as an immune-evasion factor by preventing the activation of the host’s RIG-I-mediated type I interferon (IFN) and chemokine responses (27), while the M protein efficiently suppresses NF-kappaB-dependent gene expression (28).

Together, RABV proteins influence the balance between viral replication, innate immune sensing, and immune engagement. Thus, rather than defining a simple progression from immune evasion to immune activation, different RABVs may establish distinct neuroimmune response programs depending on their combined protein and replication properties.

Although neuroimmune responses associated with infection by individual RABVs have been described, comparative analyses directly linking viral genetic variation with spatial and temporal CNS immune responses across diverse pathogenic and attenuated viruses remain limited. Differences in experimental systems, host models, inoculation routes, and analysis approaches have further complicated direct comparisons between studies. A systematic evaluation of genetically distinct RABVs under standardized conditions is therefore needed to define how viral genetic variation shapes CNS immune responses.

We investigated this question by integrating quantitative brain spatial imaging and proteomics, and analyzing field-derived, laboratory-adapted, and SAD-derived vaccine viruses under standardized conditions. An important question was how differences in neuroimmune responses among highly pathogenic RABVs relate to viral burden. More broadly, this approach allowed us to examine how the relationship between viral burden and CNS immune engagement varies among viruses during the early stages of infection.

## Results

### Genetic and functional diversity of RABV drives distinct early infection phenotypes

To investigate how genetic variation among RABVs affects neuroimmune responses in the early timepoints of the CNS infection, we used a mouse intracranial infection model with panel of recombinant viruses, including highly pathogenic (rDogA, rFox), less pathogenic (rCVS-11), and attenuated viruses (Fig. 1A, B) when peripherally administered (29, 30). The attenuated SAD L16 is neurovirulent when inoculated directly in the brain (19, 20). The further attenuated rSAD GASGAS is derived from SAD L16 with key virulence (R333E) and stabilizing (N194S) mutations in the glycoprotein, together with duplication of the glycoprotein gene (15, 18, 20).

**Figure 1.**
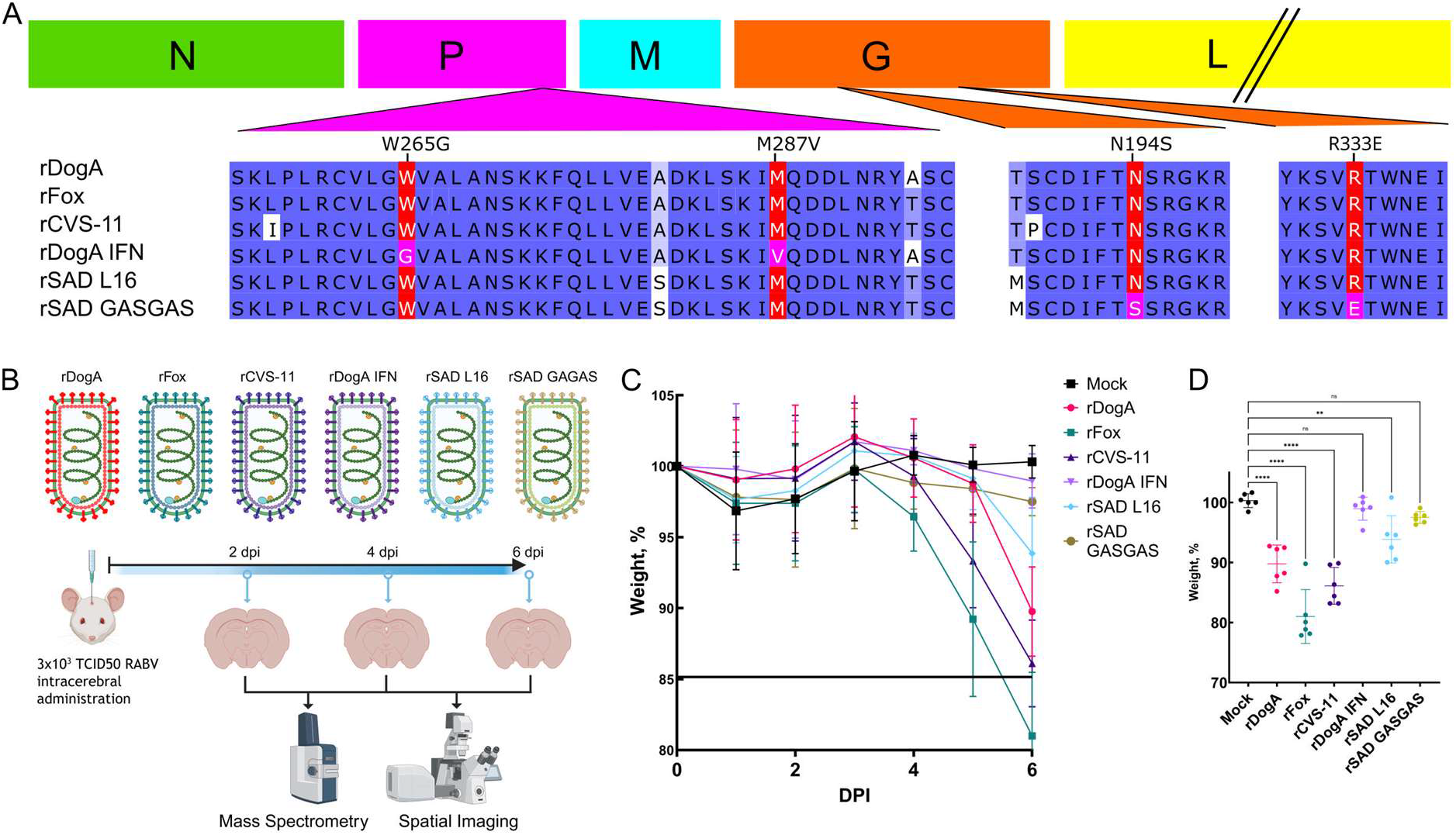
Protein sequence features of recombinant RABV and virus-specific pathogenicity in i.c. inoculated BALB/c mice. (A) RABV genome organization (N, P, M, G, L) with selected sequence amino acid alignments. The C-terminal region of P (residues 255-297) highlights rDogA IFN-specific substitutions (W265G and M287V, magenta), and the G segments (residues 187-199 and 329-338) indicate the virulence-associated residue at position 333 and the stabilizing residue at 194, both of which are substituted in rSAD GASGAS (magenta). Note that the rSAD GASGAS genome encodes two copies of G. (B) Experimental design. BALB/c mice were infected intracerebrally and analyzed at 2, 4, and 6 days post-infection (dpi) by mass spectrometry and spatial profiling by confocal microscopy following 100 µm precision sectioning. (C) Body-weight change time course. Data represent mean ± SD. (D) Body-weight change at 6 dpi. Ordinary one-way ANOVA was used to compare infected groups with mock controls. *P < 0.05, **P < 0.01, ***P < 0.001, ****P < 0.0001.

To dissect how impaired interferon antagonism influences RABV - host interactions *in vivo*, we included a recombinant rDogA (rDogA IFN) carrying the W265G and M287V substitutions in the C-terminal domain of the phosphoprotein. These mutations have previously been shown to disrupt STAT1/STAT2 binding and impair type-I interferon antagonistic activity while largely preserving viral replication capacity (31). We first assessed the replication kinetics and IFN antagonistic properties of the rDogA IFN *in vitro* (Fig. S1). The mutant virus displayed replication kinetics comparable to those of parental rDogA in BsrT7/5 cells, and reduced in Na42/13 cells (Fig. S1A, B), indicating cell type-specific influence of the introduced P protein alterations on viral propagation. Further, infection of Na42/13 cells with rDogA IFN resulted in enhanced interferon-responsive gene activation following IFN-α stimulation, consistent with reduced suppression of type-I IFN signaling (Fig. S1C). A comparable enhancement was not observed in single residue substitution (W265G or M287V) mutants. Together, these findings establish rDogA IFN as a replication-capable RABV variant with impaired interferon antagonism, providing a tool to investigate the contribution of viral immune evasion to neuroimmune regulation *in vivo* in field virus genetic background.

Variability across the viruses was primarily concentrated in the P, M, and G proteins, whereas N and L remained highly conserved (Table S1). Relative to rDogA, amino acid sequence identities for all viruses except rDogA IFN ranged from 93.32 to 95.61% for G, 91.58 to 99.33% for M, and 92.93 to 97.80% for P, while N and L proteins showed higher conservation (98.67 – 99.56% and 97.18–97.56% identity, respectively) (Table S1). These patterns of sequence conservation and divergence define the molecular landscape underlying the comparative analysis of recombinant RABVs.

We next investigated how the distinct genetic backgrounds of recombinant RABVs influence early brain infection outcomes in vivo following mouse intracerebral inoculation. Within 6 days post inoculation (dpi), the infections resulted in clear differences in disease severity across viruses (Fig. 1C, D). While no pronounced clinical signs were observed at 2 and 4 dpi, at 6 dpi virus-dependent reduction in body weight was evident. rFox-infected mice showed the most pronounced changes, retaining 81 ± 4% of their initial body weight (mean ± SD), followed by rCVS-11 (86 ± 3%), rDogA (89 ± 3%), and rSAD L16 (94 ± 4%) (Fig. 1D). Similar to the mock controls, no weight loss was observed for rSAD GASGAS and rDogA IFN.

### Temporal evolution of virus-specific neuroimmune responses

To correlate host responses with infection phenotypes, we performed time-resolved proteomic profiling of spatially defined brain tissue slices from infected animals. Principal component analysis (PCA) of proteins enriched relative to mock controls revealed a progressive emergence of virus-specific proteomic signatures over the course of infection (Fig. 2A). While enriched protein profiles showed no clear separation of RABVs at 2 dpi, partial virus clustering became apparent at 4 dpi and developed into distinct virus-specific patterns by 6 dpi, indicating progressive divergence of host response programs during infection.

**Figure 2.**
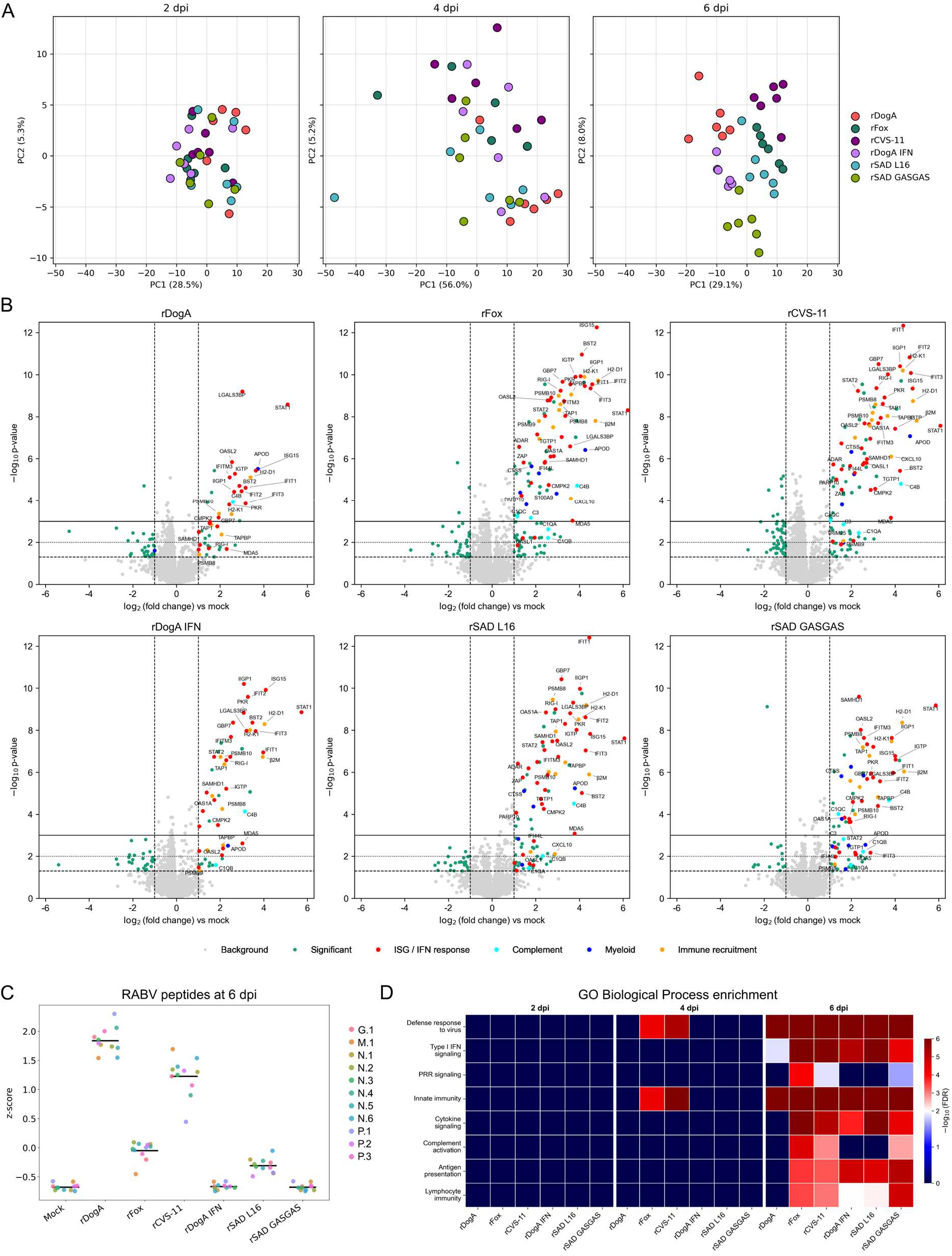
Progressive neuroimmune remodelling during RABV infection of the mouse brain. (A) PCA of proteins enriched relative to mock controls reveals increasing virus-specific divergence over time, most pronounced at 6 dpi, reflecting infection-driven remodelling of host neuroimmune pathways. (B) Volcano plots depicting differential abundance of proteins between each RABV and mock at 6 dpi. Each dot represents a protein, with highlighted significant log2 fold changes (≥ 1) and –log10 p-values (p < 0.05). (C) Z-scored abundance of detected RABV peptides in brain tissue at 6 dpi. (D) Time-resolved Gene Ontology (GO) Biological Process enrichment heatmap relative to mock summarizing immune-related pathways across viruses and time points. Color intensity indicates −log₁₀(FDR), capped to enable comparison across pathways.

At 6 dpi, a coordinated activation of antiviral and inflammatory pathways was observed across RABVs, with particularly broad and pronounced upregulation of ISGs and interferon signaling proteins, including Ifit and Oas family members, Isg15, and Stat1/2. This antiviral response was also evident in rDogA and rDogA IFN, but was less extensive and generally less pronounced than in the other RABVs (Fig. 2B). Myeloid-associated proteins, including ApoD, Ctss, and Lcp1, and complement components, including C1qa, C1qb, C1qc, C3, and C4b, were similarly enriched across most RABVs, whereas fewer proteins within these categories were altered in rDogA and rDogA IFN. Nevertheless, individual components of these responses were detected in both viruses, including ApoD and complement-associated proteins in rDogA and ApoD and C4b in rDogA IFN. The enrichment profiles further revealed virus-specific differences in the extent of myeloid, complement, and immune-cell recruitment signatures, including Cxcl10. Across the RABVs, ApoD and C4b showed particularly consistent increases relative to mock controls, with the repeated induction of ApoD suggesting activation of a shared glial stress-response program during infection.

Virus-dependent differences in RABV protein abundance were evident at 6 dpi (Fig. 2C, Table S4). Viral proteins were readily detected in most viruses, whereas no viral proteins were detected in rDogA IFN and rSAD GASGAS. Notably, both rDogA IFN and rSAD GASGAS viruses mounted detectable host proteomic responses despite the absence of measurable viral proteins.

Gene ontology (GO) biological process analysis revealed substantial differences in the extent of neuroimmune remodeling induced by individual RABVs (Fig. 2D). While rDogA infection was associated primarily with a moderate antiviral defense signature, rFox triggered broader activation of multiple immune-associated pathways, including PRR signaling, antiviral responses, complement activation, and immune recruitment programs. These findings indicate that RABVs pathogenic with similar pathogenic potential can establish distinct neuroimmune states that differ in the breadth and coordination of immune pathway activation. Notably, rFox and rCVS-11 showed earlier enrichment of antiviral defense and innate immunity, evident already at 4 dpi. At 6 dpi, this response broadened to include complement activation, antigen presentation, and lymphocyte recruitment, reflecting progressive immune engagement. The GO analysis did not show any enrichment of neuronal or synaptic processes across RABVs, indicating that the major proteomic differences were associated with immune rather than neuronal or synaptic processes.

Interestingly, rDogA, rDogA IFN, and rSAD L16 showed a more limited enrichment of pathways associated with pattern recognition receptor (PRR) signaling, complement activation, and immune recruitment compared with viruses exhibiting broader neuroimmune remodeling (Fig. 2D). However, individual proteins associated with these pathways were also altered in these viruses, indicating that reduced pathway-level enrichment reflects differences in the breadth and magnitude of the response rather than its complete absence. Distinct RABVs therefore appear to engage divergent neuroimmune programs. In contrast, rSAD GASGAS displayed broad enrichment of complement-associated pathways and the strongest immune recruitment signature, together with enrichment of PRR-associated processes, suggesting a particularly coordinated neuroimmune response characterized by innate immune activation and immune cell engagement.

### Distinct host response programs emerge from RABV genetic variation

To identify the molecular programs underlying distinct neuroimmune responses, we performed comparative proteomic analyses between the RABV with divergent infection phenotypes. Given its broad immune activation, we first focused on rFox and compared its proteomic signature with the pathogenic reference virus rDogA and the attenuated rSAD GASGAS (Fig. 3A, B). Relative to rDogA, rFox infection resulted in extensive proteomic changes, characterized by stronger induction of ISGs, myeloid-associated inflammatory proteins (S100a9, Ctss), complement components (C1qa, C1qb, C3), and antigen presentation/immune recruitment markers (H2-D1, H2-K1, Cxcl10). In contrast, comparison of rFox with rSAD GASGAS revealed a more selective divergence, with enhanced ISG expression, increased S100A9, and elevated Cxcl10 in rFox, but no significant differences in complement-associated proteins or other major immune markers. These findings indicate that rFox and rSAD GASGAS share activation of antiviral and immune recruitment programs, but differ in the extent and composition of inflammatory responses.

**Figure 3.**
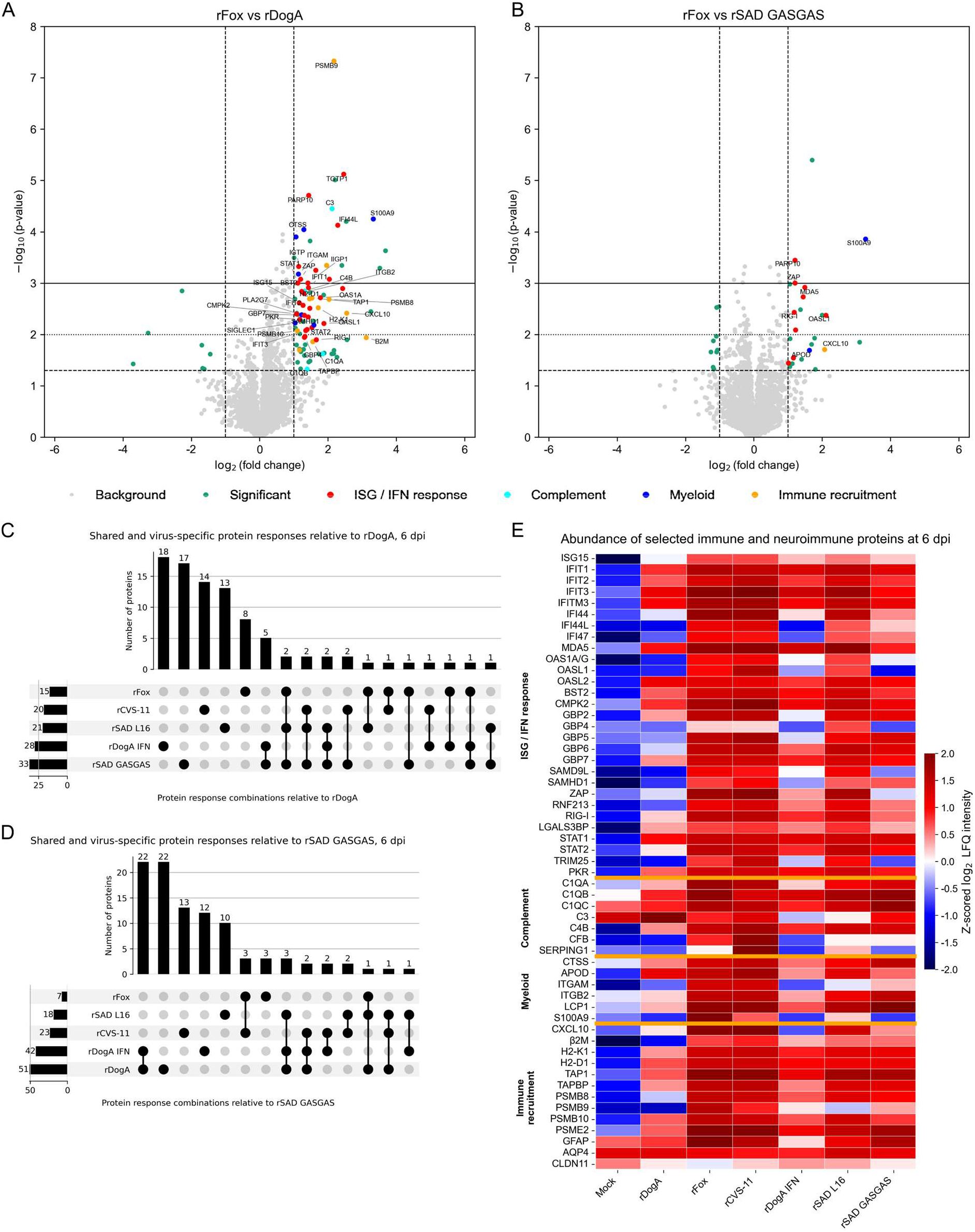
RABV virus-specific proteomic signatures reveal distinct neuroimmune responses in mouse brain. (A, B) Volcano plots depicting differential abundance of proteins between rFox and rDogA (A) and between rFox and rSAD GASGAS (B) at 6 dpi. Each dot represents a protein, with highlighted significant log_2_ fold changes (≥ 1) and –log^10^ p-values (p < 0.05). (C, D) UpSet plots summarizing unique and shared differentially abundant proteins associated with infection by individual RABV. Protein sets were generated after removal of mock-associated changes and represent protein responses relative to either rDogA (C) or rSAD GASGAS (D) reference infections. Differentially abundant proteins were defined as those with log_2_ fold change (≥ 1) and –log_10_ p-values (p < 0.05). Horizontal bars represent the number of proteins belonging to each intersection, while the dot matrix indicates the viruses contributing to each shared protein set. (E) LFQ-based heatmap of selected immune and neuroimmune proteins shown as row-wise z-scored log₂ LFQ intensities.

Similar temporal comparisons across all RABV further demonstrated that each virus established a distinct proteomic trajectory relative to the rDogA and rSAD GASGAS reference states (Fig. S4–S9), revealing both shared and virus-specific patterns of host response. To define the extent of overlap between these responses, datasets were first corrected by excluding enriched proteins detected in the corresponding mock-infected controls, followed by UpSet analysis using rDogA and rSAD GASGAS as reference backgrounds (Fig. 3C, D). Beyond immune-associated pathways, rFox-specific proteins relative to rDogA included regulators of mitochondrial respiratory chain assembly (Coa6, Surf1), endoplasmic reticulum and membrane homeostasis (Dpm1, Pgap1, Reep1), and cellular stress adaptation (Paox, Nedd4l), suggesting that rFox infection imposes additional metabolic and cellular stress on brain tissue. In contrast, comparison of rFox with rSAD GASGAS identified only a limited set of differentially abundant proteins, including Osgep, Slc7a6os, and Ubxn7, suggesting substantial overlap in the antiviral and immune-associated proteomic responses induced by these highly pathogenic and highly attenuated viruses, with remaining differences primarily involving cellular metabolism and protein homeostasis.

Further analyses of the different RABVs relative to the rDogA and rSAD GASGAS references identified common infection-associated proteins as well as virus-specific signatures, highlighting distinct molecular programs associated with different RABV-induced neuroimmune states (Fig. 3C, D, S4–S9). Interestingly, although rDogA IFN infection induced a pronounced antiviral proteomic response relative to mock controls, direct comparison with parental rDogA revealed only a limited set of differentially abundant proteins. This suggests substantial convergence between the two infection states, with the mutations primarily enhancing the efficiency of an existing antiviral program rather than establishing a distinct inflammatory state.

**Figure 4.**
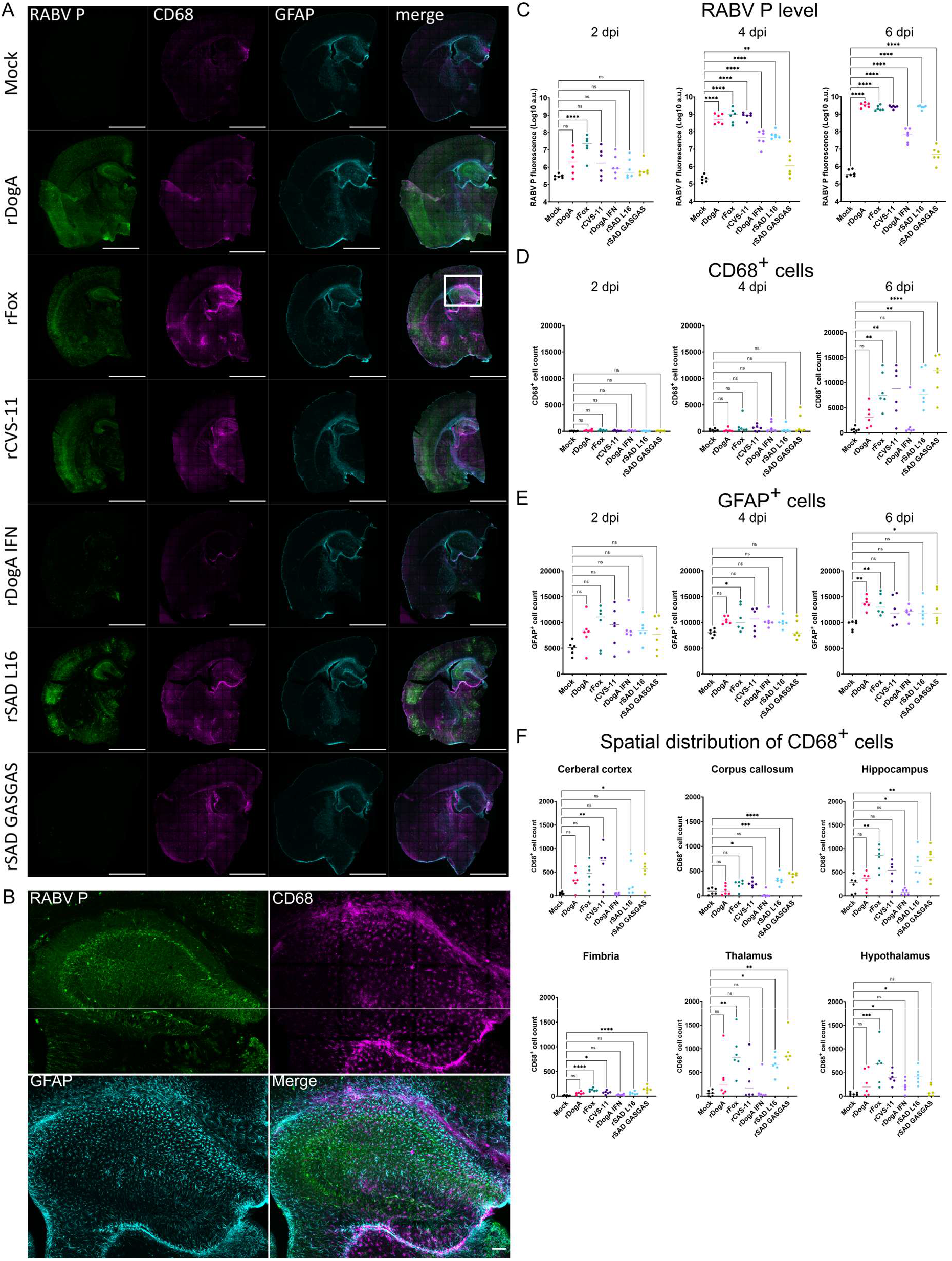
RABV establish distinct spatial patterns of infection and neuroimmune activation. (A) Confocal imaging of brain infection and neuroimmune response at 6 dpi. Representative images of RABV P (green), CD68 (magenta) and GFAP (cyan) from 6 replicate mice for all viruses and mock control are shown. Scale bar 2 mm. White square selection in rFox merged image indicates a region with detailed view in (B) showing individual infected cells, CD68^+^ cells and GFAP^+^ astrocytes, scale bar 100 µm. (C – E) Quantitative analysis of RABV P (C), CD68^+^ cell (D) and GFAP^+^ cell levels (E) at 2, 4 and 6 dpi. (F) Spatial distribution of CD68^+^ cells in quantified brain regions at 6 dpi. Points corresponding to individual animals and median are shown. Ordinary one-way ANOVA was used to compare infected groups with mock controls. *P < 0.05, **P < 0.01, ***P < 0.001, ****P < 0.0001.

Comparison of rSAD L16 and rSAD GASGAS identified a limited set of virus-specific proteins, which were primarily associated with immune-related processes, cellular organization, and stress adaptation rather than canonical antiviral pathways. Notably, rSAD L16-specific enrichment of CD34 and Pla2g7 suggested altered immune - tissue interactions and inflammatory lipid metabolism. In addition, proteins involved in intracellular trafficking and structural organization, including Ap3s2, Jakmip1, and Specc1, pointed toward virus-dependent differences in vesicular transport and cytoskeletal regulation. A further subset of proteins associated with protein handling and cellular homeostasis, including Bag4, Pfdn6, Nat14, and Rps28, suggested altered regulation of protein folding, translation, and stress adaptation. Together, these findings indicate that the divergent neuroimmune phenotypes of rSAD L16 and its variant rSAD GASGAS are not primarily distinguished by the magnitude of canonical antiviral signaling, but by distinct programs of immune-associated responses and cellular adaptation.

Consistent with the pathway-level findings (Fig. 2D), the protein-level analysis revealed virus-dependent induction of ISGs and neuroimmune responses (Fig. 3E). rFox and rCVS-11 showed the strongest immune-associated changes at 6 dpi, with earlier proteomic divergence already evident at 4 dpi in the corresponding virus comparisons (Fig. S5, S8), indicating accelerated activation of host response programs in these infections. Focused analysis of immune and neuroimmune proteins further identified altered abundance of specific host-defense components in rSAD GASGAS consistent with its distinct neuroimmune state (Fig. 3E).

Canonical astrocyte markers such as Gfap and Aqp4 (Fig. 3E), as well as the tight junction protein Cldn11, were detected in the proteomic datasets but did not show differential abundance in comparisons to either rDogA or rSAD GASGAS indicating no broad change in these canonical astrocyte-associated proteins at the proteome level.

Collectively, these findings indicate that RABVs elicit qualitatively distinct neuroimmune responses, characterized by differences in antiviral signaling, inflammatory responses, immune-associated proteins, and cellular adaptation in ways that cannot be explained solely by differences in viral accumulation.

### Spatial uncoupling of viral dissemination and CNS immune activation

To define the spatial and cellular context of infection and immune activation, we performed immunofluorescence analysis of RABV P, CD68 (microglia/macrophages), and GFAP (astrocytes) levels across time points and brain regions (Fig. 4, S10–S14).

At 2 dpi, RABV P signal was low across most viruses, with more pronounced infection detected in rFox, suggesting faster virus propagation (Fig. 4C, Fig. S10). However, no associated CD68 response was present (Fig. 4D). Similarly, GFAP levels remained low and comparable to mock (Fig. 4E).

By 4 dpi, viral antigen levels strongly increased in all viruses, with rDogA and rCVS-11 reaching a level similar to rFox. Markedly, in rDogA IFN and rSAD L16 the increase was significantly lower, and in rSAD GASGAS only a modest presence of RABV P was observed (Fig. 4C). No substantial CD68^+^ cell accumulation was detected (Fig. 4D, E; Fig. S11). GFAP levels in astrocytes increased further in both infected and mock animals to a comparable extent, with only rFox showing a modestly higher elevation (Fig. 4E). This indicates that early infection does not trigger a full-scale immune response, but rather leads to limited host perturbation in viruses with faster infection kinetics.

To assess regional differences in viral distribution, RABV P abundance was quantified across six brain regions, including the cerebral cortex, corpus callosum, hippocampus, fimbria, thalamus, and hypothalamus (Fig. S12-S14). Spatial analysis revealed that rFox was already detectable in all examined brain regions at 2 dpi, indicating early widespread CNS dissemination. Similarly, for viruses with detectable viral antigen at later time points, RABV P was observed across all analyzed regions, with no consistent regional preference detected.

A pronounced divergence between viruses emerged at 6 dpi. High levels of viral antigen were detected in rDogA-, rFox-, and rCVS-11-infected brains. Consistent with previous reports (30), rSAD L16 also showed substantial RABV P levels comparable to those observed in highly pathogenic viruses. In contrast, rDogA IFN and rSAD GASGAS maintained markedly reduced viral antigen levels, similar to those observed at 4 dpi, suggesting limited viral dissemination or accumulation in these infections (Fig. 4 A-C).

Interestingly, spatial analysis revealed a delayed pattern of hypothalamic infection for rSAD GASGAS, with lower RABV P signal at 4 dpi than in other viruses, followed by comparable levels of infection at 6 dpi (Fig. S14).

Notably, CD68^+^ cells increased sharply at 6 dpi in responses induced by rSAD GASGAS, rSAD L16, rCVS-11, and rFox (Fig. 4 A,B), while no significant response to rDogA and rDogA IFN was detected (Fig. 4A,D). This indicates that immune activation in the CNS is highly virus-dependent, with certain viruses inducing stronger CD68^+^ myeloid response, regardless of viral burden.

In contrast to the myeloid response, GFAP⁺ astrocyte responses exhibited a more gradual increase over time. At 6 dpi, the strongest increase in GFAP⁺ cell numbers was observed following infection with rDogA and rFox, whereas rSAD GASGAS showed only a modest elevation with substantial inter-animal variability. No significant increase was observed for the remaining viruses. This suggests that while astrocytes respond early in infection, their activation intensity is delayed and more sustained in certain viruses, correlating with the escalation of overall immune activation. Notably, these spatial changes in GFAP⁺ cell numbers were not accompanied by significant changes in bulk GFAP protein abundance in the proteomic analysis, highlighting the complementary nature of the two approaches.

Regional analysis (Fig. 4F, S15-16) highlighted marked spatial heterogeneity in immune activation. Notably, CD68⁺ myeloid cells were enriched in response to rFox in several regions including the hippocampus, fimbria, thalamus and hypothalamus. rSAD GASGAS showed a similar myeloid response in hippocampus, fimbria, thalamus, and additionally in corpus callosum, but no detectable increase in the hypothalamus (Fig. 4F). Notably, this absence of a hypothalamic myeloid response persisted in spite of the comparable levels of RABV P detected in this region at 6 dpi (Fig. S14).

Notably, the appearance of CD68 signal was not spatially colocalized with infection (Fig. 4A, B), indicating that local immune activation did not simply mirror the distribution of viral antigen. Instead, immune activation was concentrated in specific brain regions, with the hippocampus, fimbria and thalamus exhibiting stronger responses. GFAP⁺ astrocytes showed a more diffuse profile with higher cell numbers in the hippocampus and thalamus to rFox and rSAD GASGAS, and lower cell numbers in the cortex (Fig. S15–16), supporting the idea that neuroimmune responses are regionally heterogeneous and vary between viral infections.

Together, these data reveal that RABV infection triggers a temporally and spatially distinct neuroimmune response, where immune activation in the CNS varies between viruses and is spatially and temporally distinct. Highly pathogenic rFox combines strong infection with pronounced immune activation, whereas attenuated rSAD GASGAS also elicits strong immune responses, but with lower viral loads, highlighting the dynamic interplay between viral burden, immune engagement, and shaping the course of infection.

### Complement C3 responses reveal distinct cellular modes of neuroimmune activation

To further define neuroimmune activation at the level of complement signaling, we assessed the spatial distribution of complement component C3 in relation to viral infection and astrocyte activation at 6 dpi (Fig. 5, Fig. S17). Confocal imaging revealed a widespread C3 signal across multiple brain regions in infected animals in most RABV viruses (Fig. 5A). C3 signal was frequently associated with GFAP⁺ astrocytes, consistent with previous reports identifying C3 as a marker of a distinct reactive astrocyte state (32). Quantitative analysis confirmed strong increase in total C3 level in rFox, rCVS-11, rSAD L16 and rSAD GASGAS, whereas rDogA and rDogA IFN groups did not show significant differences compared with mock controls, despite some inter-animal variability in the rDogA. (Fig. 5B).

**Figure 5.**
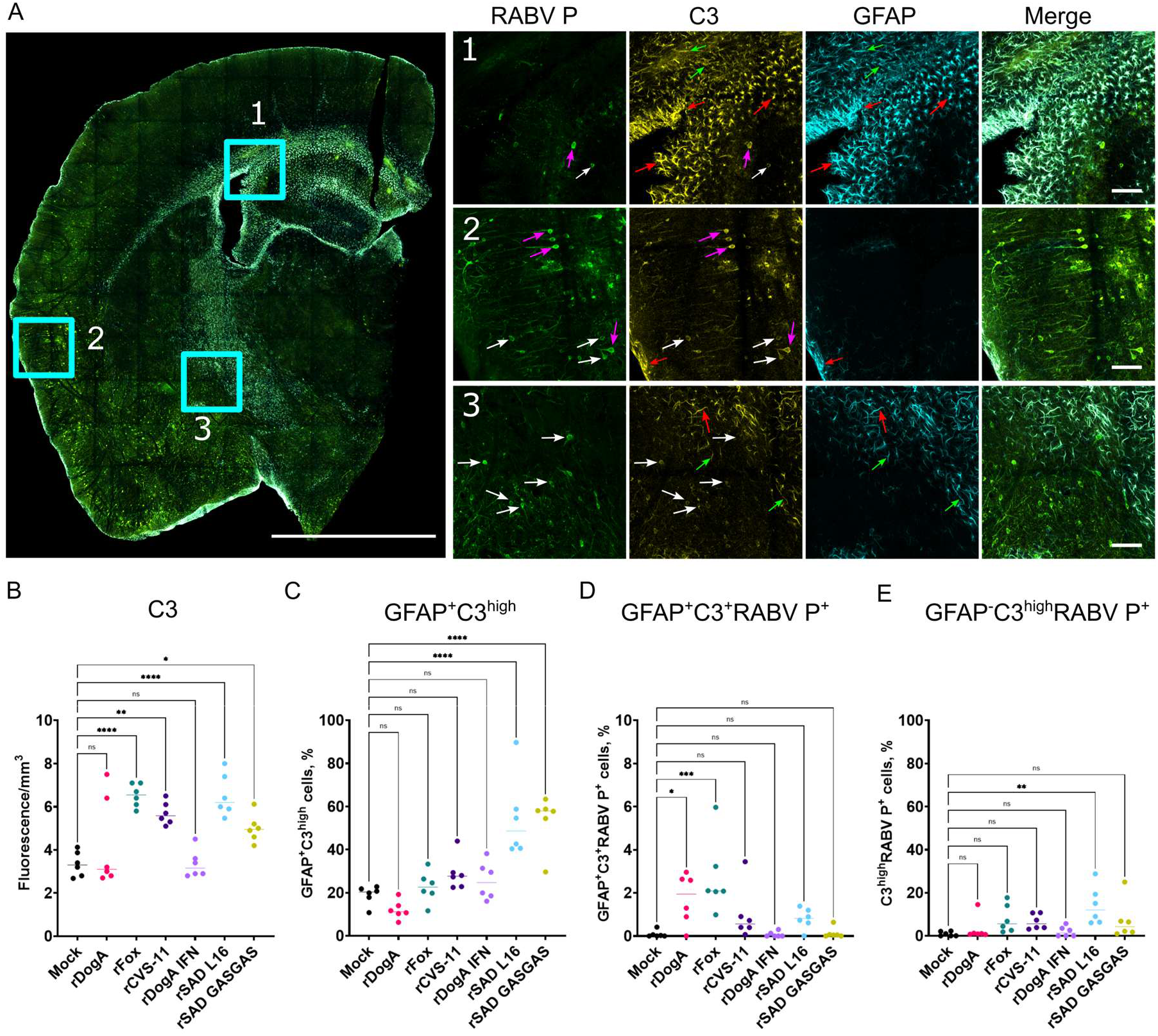
RABV infection drives spatially organized complement C3 activation in the brain. (A) Whole brain section view of RABV P (green), C3 (yellow) and GFAP (cyan) at 6 dpi. Representative image following rCVS-11 infection. Scale bar 2 mm. Square selections indicate regions with detailed views on the right (1, 2, 3). The arrows indicate examples of C3^high^GFAP^+^ (red), C3^low^GFAP^+^ (green), C3^high^GFAP^-^RABV P^+^ (magenta) and C3^low^GFAP^-^RABV P^+^ (white) cells. Scale bar 100 µm. (B) C3 fluorescence levels across whole brain sections. (C, D, E) Fractions of GFAP^+^C3^high^ (C), GFAP^+^C3^high^RABV P^+^ (D) and GFAP^-^C3^high^RABV P^+^ (E) cells. Points corresponding to individual animals and median are shown. Ordinary one-way ANOVA was used to compare infected groups with mock controls. *P < 0.05, **P < 0.01, ***P < 0.001, ****P < 0.0001.

We noted the presence of cells with low and high C3 levels, and abundant colocalization of C3 with GFAP^+^ cells (Fig. 5A) suggesting that astrocytes may be a significant cellular source of C3 upregulation. To quantitively assess it, we determined the fractions of C3^high^GFAP^+^ and C3^high^GFAP^-^ cells (Fig. 5A). Strikingly, such targeted analysis revealed a pronounced increase in C3^high^ proportion in GFAP⁺ astrocytes only in rSAD L16 and rSAD GASGAS (Fig. 5C). Notably, this astrocyte-associated C3 response did not simply correlate with infection. Although it was detectable for rDogA and rFox, only a small fraction of C3^+^ astrocytes was infected, whereas infection of this cell population was minimal or absent for other viruses (Fig. 5D). Thus, direct astrocyte infection alone is unlikely to account for the virus-specific increase in GFAP^+^C3^high^ cells and may instead contribute to differential modulation of astrocyte responses.

Although rFox and rCVS-11 infection induced widespread C3 accumulation throughout the brain, this increase was not reflected by elevated C3^high^GFAP^+^ astrocytes in these viruses, suggesting that C3 induction during pathogenic infection involves additional cellular sources. C3 was also detected in GFAP⁻ cells, including RABV P⁺ cells. However, the fraction of C3^high^GFAP⁻ cells was significantly increased only following rSAD L16 infection and remained low with the other viruses (Fig. 5E). Thus, the widespread increase in C3 observed with rFox and rCVS-11 appears to reflect increased C3 across a broader population of cells rather than expansion of a distinct C3^high^ cellular population. These findings reveal distinct cellular patterns of complement-associated neuroimmune responses shaped by virus identity.

### Adaptive immune recruitment is uncoupled from viral burden

To investigate T cell recruitment, we performed immunofluorescence analysis of RABV P, CD3, and CD45 across multiple viruses and time points (Fig. 6).

**Figure 6.**
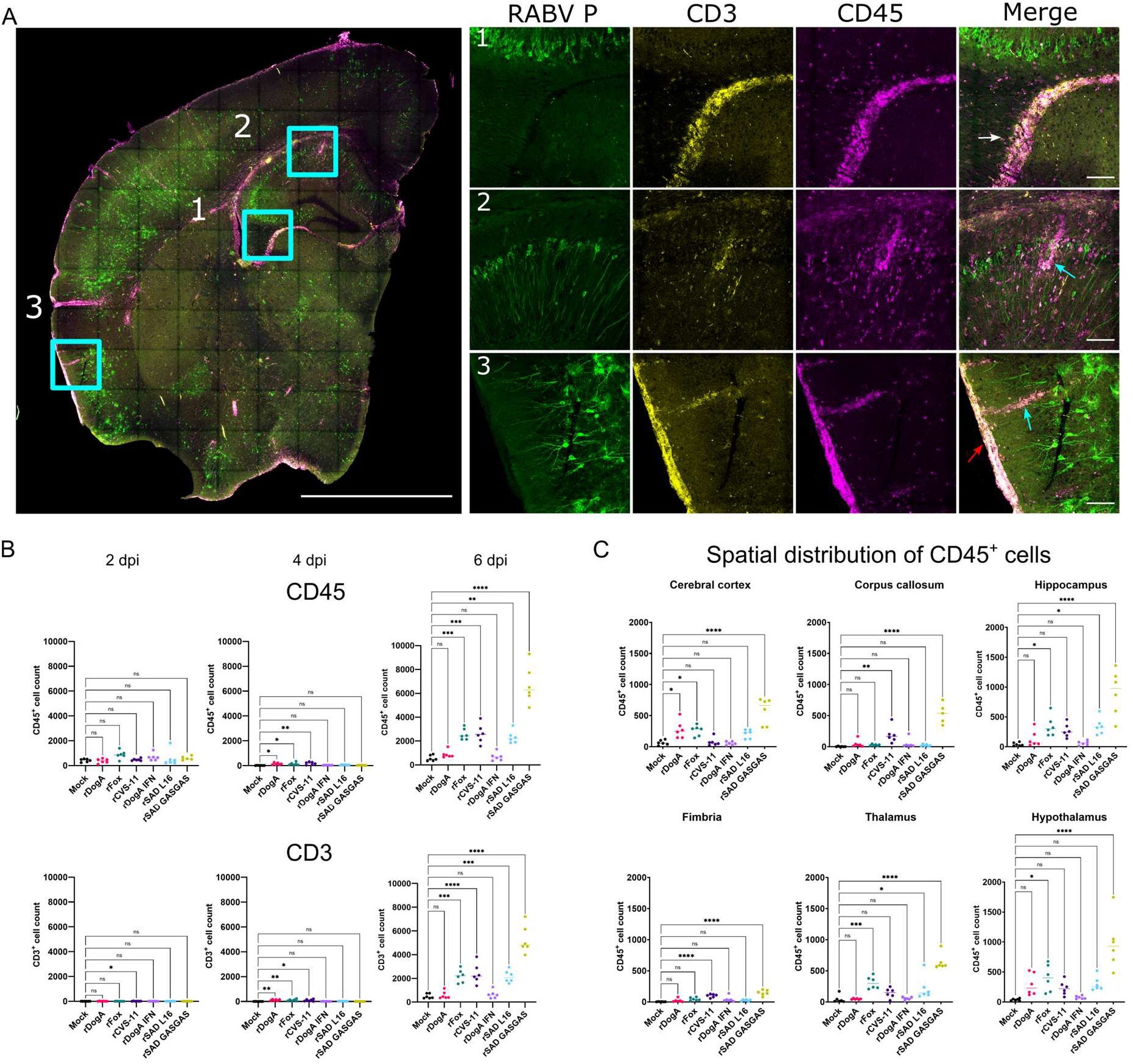
RABV induce distinct spatial patterns of T cell recruitment in the brain. (A) Whole brain section view of RABV P (green), CD3 (yellow) and CD45 (magenta) at 6 dpi. Representative image following rSAD L16 infection. Scale bar 2 mm. Square selections indicate regions with detailed views on the right (1, 2, 3) showing strong T cell infiltration near ventricular system, vascular interfaces, and brain periphery indicated by white, cyan and red arrows, respectively. 1 - region encompassing ventricle, thalamus and fimbria, 2 – region encompassing hippocampus, ventricle and corpus callosum, 3 – region at the brain periphery. Scale bar 100 µm. (B) Quantification CD45 and CD3 fluorescence across whole brain sections. (C) Spatial distribution of CD45^+^ cells in brain regions at 6 dpi. Strongest response in hypothalamus, hippocampus and cerebral cortex. Points corresponding to individual animals and median are shown. Ordinary one-way ANOVA was used to compare infected groups with mock controls. *P < 0.05, **P < 0.01, ***P < 0.001, ****P < 0.0001.

At 2 and 4 dpi, minimal CD45⁺ and CD3⁺ cell infiltration was observed in all viruses, consistent with limited immune cell engagement during the early stages of infection (Fig. 6B; Fig. S18–S20). Only minor fluctuations were observed, indicating that early host responses are dominated by innate mechanisms.

By 6 dpi, robust lymphocyte infiltration was observed in a virus-dependent manner. Strong increases in CD45⁺ and CD3⁺ cells were detected in rFox-, rCVS-11-, rSAD L16-, and particularly rSAD GASGAS-infected brains, whereas rDogA and rDogA IFN showed little to no recruitment and remained comparable to mock controls (Fig. 6B). Notably, rSAD GASGAS, even with its low viral burden, induced the most pronounced T cell response, highlighting the strong capacity of this highly attenuated SAD L16 variant to induce CNS T-cell recruitment.

Spatial analysis revealed that infiltrating lymphocytes were distributed throughout the brain and were not directly linked to the localization of infected cells (Fig. 6A, C). Lymphocyte accumulation was prominent along the ventricular system (Fig. 6A, insets 1 and 2) and at the brain periphery (Fig. 6A, inset 3), consistent with possible recruitment through cerebrospinal fluid and meningeal interfaces. In addition, high lymphocyte accumulations were observed around structures morphologically consistent with blood vessels (Fig. 6A, insets 2 and 3), consistent with a possible association between T cells and vascular interfaces.

These data demonstrate that T cell entry to the brain is a late event during RABV infection, occurring predominantly at 6 dpi, and strongly influenced by specific viruses. Importantly, the vaccine virus rSAD GASGAS elicits the strongest lymphocyte infiltration, whereas rDogA and rDogA IFN showed little to no recruitment. This indicates that the magnitude of CNS T cell recruitment is not simply determined by viral burden, but depends on virus-specific host interactions.

## Discussion

By analyzing a panel of genetically distinct RABV within the same experimental framework, this study demonstrates that viral genetic variation shapes distinct neuroimmune profiles, characterized by differences in viral accumulation, antiviral signaling, complement activation, glial responses, and immune engagement. This is particularly relevant in the CNS, where neurotropic viruses must balance replication with immune control and insufficient antiviral responses can permit viral spread, whereas excessive immune activation may contribute to neurological pathology (1–3). However, our data reveal that, following direct CNS infection, the patterns of neuroimmune activation are not uniform among RABV with high pathogenic potential. Thus, the consequences of RABV infection in the CNS appear to depend on how viral accumulation and host immune responses are coordinated rather than on the magnitude of either alone, shaping distinct neuroimmune states (Fig. 7). High viral accumulation in rDogA was associated with comparatively limited immune engagement, whereas rSAD GASGAS elicited pronounced immune responses even though viral levels were minimal. rDogA IFN further illustrated that enhanced antiviral signaling can restrict viral accumulation without triggering broad secondary immune activation. Conversely, rFox, rCVS-11, and rSAD L16 combined substantial viral burden with pronounced immune engagement.

**Figure 7.**
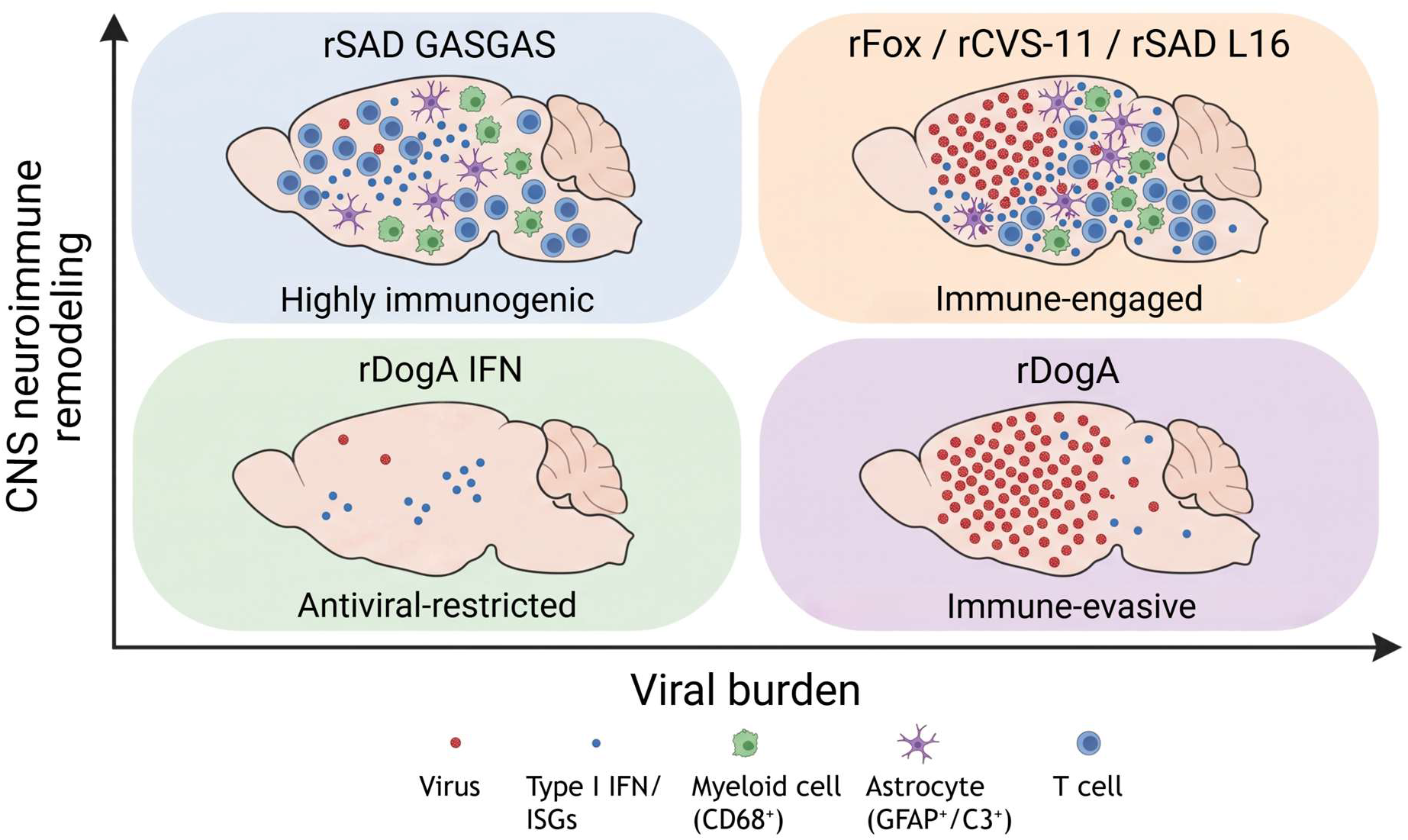
Distinct neuroimmune states during RABV infection. Conceptual model summarizing how genetically distinct RABVs establish different relationships between viral burden and CNS immune remodeling. Viruses are positioned schematically according to their relative viral antigen abundance and extent of neuroimmune activation observed in proteomic and spatial imaging analyses. rDogA represents an immune-evasive, virus-dominant state characterized by high viral burden and limited CNS neuroimmune remodeling, whereas rDogA IFN represents an antiviral-restricted state with reduced viral accumulation, enhanced antiviral signaling, and limited neuroimmune remodeling. rFox, rCVS-11, and rSAD L16 occupy an immune-engaged state characterized by substantial viral burden together with coordinated CNS immune responses. In contrast, rSAD GASGAS represents a highly immunogenic, low-virus state characterized by strong CNS immune engagement despite minimal detectable viral protein.

One striking observation is the marked divergence between the two pathogenic wild-type viruses rDogA and rFox. Albeit both retain high pathogenic potential and the capacity to cause lethal disease following peripheral infection (29), they establish distinct neuroimmune states in the CNS (Fig. 7). rDogA infection was associated with substantial viral accumulation but comparatively limited complement activation and myeloid engagement, whereas rFox induced early and broad activation of antiviral, complement-, chemokine-, and myeloid-associated pathways. This contrast demonstrates that high pathogenic potential can be associated with markedly different CNS immune profiles and levels of activation, consistent with distinct patterns of innate immune response evasion reported for pathogenic and attenuated RABV (6). Severe rabies CNS infection in humans and experimentally infected mice has likewise been reported in the absence of prominent inflammatory responses (33, 34). The divergent responses of rDogA and rFox therefore suggest that, even among highly pathogenic RABV, viral burden can be coupled to neuroimmune activation in different ways.

This raises the question of which viral determinants shape these neuroimmune trajectories. The mutant viruses provide experimental evidence that defined changes in the P or G proteins can redirect neuroimmune responses, whereas the differences between naturally divergent viruses represent phenotypes whose underlying genetic determinants remain unresolved. In this context, rFox, rCVS-11, and rSAD L16 induced robust immune responses characterized by strong ISG induction, complement activation (C1qa, C1qb, C1qc, C3, C4b), chemokine upregulation (Cxcl10), and pronounced CD68-associated myeloid responses, consistent with previous observations (6, 22, 35–37). In contrast, targeted modifications in viral proteins produce distinct host response phenotypes. The phosphoprotein substitutions in rDogA IFN (W265G and M287V) impair STAT1/STAT2 antagonism (31), resulting in enhanced type I interferon signaling and a distinct antiviral profile. Thus, altered P-mediated interferon antagonism can redirect the balance between viral replication and subsequent CNS immune engagement.

Glycoprotein-associated modifications likewise demonstrate that robust immune engagement does not require high viral protein accumulation. The R333E and N194S substitutions together with duplication of the glycoprotein gene in rSAD GASGAS (15, 18) were associated with pronounced immune responses and substantial T cell infiltration. Mock-corrected comparison of rSAD GASGAS and rSAD L16 identified only a limited number of modulated proteins. These shared alterations suggested conserved changes associated with immune-associated signaling (Tmem126a and Fermt3) (38, 39), intracellular trafficking and membrane dynamics (Iqsec1 and Gpcpd1)(40, 41), cytoskeletal organization (Ap3s2, Jakmip1, Specc1, and Pfdn6)(42–45), lipid metabolism and inflammatory regulation (Pla2g7)(46). The lower level of CD34 in rSAD GASGAS may reflect altered endothelial or perivascular-associated cellular states, potentially influencing vascular - immune communication and leukocyte trafficking (47, 48). Although these differences were comparatively subtle, they indicate distinct cellular adaptations that may contribute to the enhanced immune engagement of rSAD GASGAS.

An intriguing possibility is that differences in neuroimmune responses induced by RABV variants reflect, at least in part, evolutionary divergence associated with adaptation to distinct reservoir environments. Rabies virus evolution is shaped by host-specific transmission ecology, and recent work has suggested that adaptation to different reservoir species may involve changes in viral glycoprotein properties affecting neuronal receptor function and potentially host behaviour (49). Such findings raise the possibility that evolutionary pressures associated with transmission and host adaptation extend beyond viral replication and neuroinvasion to influence broader virus - host interactions within the CNS. In this context, the distinct neuroimmune profiles induced by rDogA and rFox may reflect differences in their evolutionary histories, although the contribution of host adaptation to these phenotypes remains to be established.

Our findings extend previous studies showing that viral genetic determinants, particularly those affecting interferon antagonism and replication capacity, influence CNS immune activation and disease outcome in individual RABV variants (6, 10, 14, 18, 30, 36, 37, 50–56). Recent high-dimensional approaches, including single-cell transcriptomics, have further identified virus-associated differences in CNS immune cell states and transcriptional programs (35, 57). However, these studies have primarily focused on cellular identities and molecular signatures, leaving the temporal, proteome-wide, spatial, and coordinated organization of neuroimmune responses across genetically distinct RABV insufficiently characterized.

Although bulk proteomics and spatial immunofluorescence provided broadly consistent estimates of viral abundance, with both approaches identifying rDogA IFN and rSAD GASGAS as low-burden infections, differences were also apparent in the relative ranking of the remaining viruses. These measurements capture related but distinct aspects of viral protein accumulation. Proteomic analysis reflects the abundance of detectable viral peptides across the bulk tissue sample, whereas immunofluorescence specifically measures the spatial distribution and abundance of antibody-detectable RABV P protein in the examined sections. Thus, the two approaches provide complementary rather than directly interchangeable measures of viral abundance.

Beyond virus-specific differences in antiviral and immune activation, astrocyte-associated and complement-related changes emerged as another major component of the virus-dependent response. These neuroimmune responses were not uniform at the cellular level, with complement- and astrocyte-associated responses providing one example of how similar levels of overall immune engagement can arise through different cellular programs. C3 showed increased abundance and preferential colocalization with astrocytes, supporting an astrocyte-associated complement response. In contrast, canonical astrocyte markers such as Gfap and Aqp4 were not differentially abundant, while Cldn11, a tight-junction protein and major component of CNS myelin (58), was likewise unchanged. Thus, the observed changes point to selective modulation of astrocyte-associated complement and immune communication rather than broad changes in astrocyte marker expression. RABV infection induces both shared and virus-specific astrocyte-associated responses involving complement regulation and immune communication, consistent with emerging evidence that astrocytes form dynamic, regionally organized networks capable of functional adaptation (59). ApoD was consistently upregulated across all tested RABV, indicating a conserved lipid- and stress-associated CNS response. Recent work identified ApoD as a potential proviral host factor that interacts with RABV G and promotes cholesterol-associated viral propagation, suggesting that ApoD induction may reflect both host metabolic adaptation and virus-supportive effects (60). However, ApoD has also been associated with microglial transcriptional responses to RABV infection (57) and the bulk proteomic data presented here cannot establish the cellular origin of this response. Its induction may therefore reflect astrocyte-associated, microglial, or broader multicellular remodeling.

Previous studies have shown that astrocyte infection is strongly dependent on virus and route of infection, with differences in astrocyte tropism observed among RABV variants (30). Astrocytes can also mount cell-intrinsic antiviral and inflammatory responses to RABV through RIG-I/MDA5 - MAVS signaling, with attenuated viruses inducing stronger innate responses than wild-type virus (61). More recent studies in human brain cell models have identified distinct astrocyte states associated with RABV infection and a pre-existing pro-viral astrocyte subpopulation (62). Moreover, defined glycoprotein mutations have recently been shown to alter astrocyte tropism and protective CNS immunity in vivo (51). These studies and our data point to an important role for astrocytes in RABV - host interactions. Although the bulk proteomic and imaging approaches used here do not resolve individual astrocyte subpopulations, the differential abundance of astrocyte-associated immune proteins suggests that viral genetic variation influences how RABV engages astrocyte-associated functions. Importantly, astrocyte- and complement-associated responses were not uniformly coupled to interferon activation. rDogA IFN, for example, displayed enhanced antiviral signaling without a corresponding astrocyte-associated response, whereas other viruses showed coordinated activation of interferon-, complement-, and glial-associated pathways. This dissociation further supports the concept that different RABV establish qualitatively distinct neuroimmune states rather than merely different intensities of a common inflammatory program.

This study was limited to seven to eight week old BALB/c male mice, and it can be expected that the neuroimmune response profile will be affected by the strain, gender and age, which will require further investigation. Additionally, only intracerebral route of infection was used, as peripheral infection would prevent some of the viruses from reaching the CNS. However, together, our data indicate that viral genetic variation can alter how viral accumulation, antiviral restriction, and CNS immune responses are linked, generating distinct neuroimmune states. Defined changes in P and G can finely modulate these responses, whereas the phenotypes of naturally divergent viruses likely reflect the combined effects of multiple viral factors. Importantly, the highly immunogenic neuroimmune profile induced by rSAD GASGAS illustrates that vaccine-associated genetic modifications can profoundly reshape CNS responses to infection, beyond their effects on the classical attenuation phenotype.

## Methods

### Viruses

Recombinant rabies viruses rRABV DogA and rRABV Fox, derived from highly virulent field isolates, rCVS-11, derived from Challenge Virus Street-11, and rSAD L16, derived from the attenuated live vaccine virus SAD B19 (63), have been described previously (16, 29, 30) (Table S2). SAD GFP is a previously described GFP-expressing SAD L16 variant (64).

Amino acid substitutions in the phosphoprotein open reading frame (P ORF) of rDogA, namely W265G, M287V, and W265G/M287V, were introduced into the full-length cDNA clone by PCR-based site-directed mutagenesis. The single mutants W265G and M287V were generated by PCR amplification of the complete full-length cDNA plasmid using the primer pairs 1/2 and 3/4, respectively. The double-mutant plasmid prDogA P W265G/M287V (rDogA IFN), was generated by a second round of mutagenesis using the primer pair 1/2 and prDogA P M287V as PCR template. The complete viral cDNA sequences were verified by sequencing. Recombinant viruses were rescued from cDNA and subsequently amplified on NA42/13 cells as described previously (29).

For the construction of the full-length cDNA plasmid prSAD GASGAS, the synthetic DNA fragments synthGASGAS-1 (1.0 kb) and synthGASGAS-2 (1.2 kb) were inserted into the EcoRI-linearized cloning vector pBluescript SKII by Hot Fusion cloning (65). This resulted in two sequential glycoprotein (G) coding sequences separated by N/P gene-border-derived transcription signals. PCR fragments corresponding to the cloned G sequences were amplified using primer 5/6 (1.0 kb) and 7/8 (1.2 kb). These fragments were fused with a 14.4 kb DNA fragment amplified from a SAD L16 cDNA clone (66). To delete the pseudogene region, the resulting plasmid was used as template for PCR amplification of 15 kb and 1.7 kb DNA fragments using the primer pairs 9/10 and 7/11, respectively. The fragments were subsequently assembled by Hot Fusion cloning. The resulting plasmid prSAD GASGAS, containing two copies of the G gene with mutations at amino acid positions 194 and 333, was rescued in BSR T7/5 cells according to a standard protocol (66).

### Sequence alignment and identity analysis

Genbank complete RABV genomes were used for sequence alignment: LN879480.1 (DogA), LN879481.2 (Fox), M31046 (SAD B19 sequence used for SAD L16 alignment), LT839616.1 (CVS-11). Multiple sequence alignments were performed using Jalview (version 2.11.5.1) (67). Protein sequences were imported in FASTA format and aligned using the built-in alignment interface. Alignments were manually inspected and refined to ensure correct residue positioning in conserved regions.

Pairwise and global sequence identity scores were calculated within Jalview using its internal identity calculation tools, based on aligned positions and excluding gaps where appropriate. Identity values were used to assess similarity between viral protein variants and reference sequences.

Sequence snippets corresponding to regions of interest were generated directly from the alignment view by selecting aligned segments and exporting them as formatted svg files. These snippets preserve alignment spacing and residue conservation patterns, allowing clear visualization of motif-level differences between sequences.

### Animal experiments

Seven to eight week old male BALB/c mice obtained from a commercial breeder (Charles River, Germany) were used. The use of a single gender was a deliberate design choice intended to maintain experimental consistency when comparing multiple recombinant RABVs across different time points and analytical approaches. Animals were randomly assigned to experimental groups and housed in individually enriched cages with water and standard rodent chow provided ad libitum. For each rabies virus construct (Suppl. Table 1, 2), six mice were inoculated intracerebrally (i.c.) with 3 × 10³ TCID50 in a volume of 0.02 mL and humanely euthanized at 2, 4, or 6 days post infection (dpi). Mock-infected control animals (n = 6) received 20 µL PBS. The experiments were repeated independently three times to generate three replicates.

Prior to infection, mice were briefly anesthetized with isoflurane (Isofluran CP, CP Pharma, Germany) until loss of righting reflex and absence of response to tactile stimulation. Animals were manually restrained, and the injection site was disinfected. Inoculation was performed using a sterile syringe and needle, targeting the forebrain region based on anatomical landmarks between the eyes and snout to ensure consistent delivery. Following injection, mice were returned to their cages and monitored daily for body weight and clinical signs using a previously described clinical scoring system (68). To ensure synchronized sampling, animals were inoculated sequentially so that all designated groups could be euthanized on the same day according to the experimental schedule..

For terminal procedures, mice were anesthetized using a combination of isoflurane followed by ketamine–xylazine (Serumwerk Bernburg AG, Bernburg, Germany - 100 mg/kg Ketamin, 10 mg/kg) administered intraperitoneally. Adequate anesthesia was confirmed by the absence of spinal reflex responses. Animals were then secured, and thoracotomy was performed to access the heart. For transcardiac perfusion, the right ventricle was incised to allow blood outflow, and a cannula was inserted into the left ventricle. Mice were perfused with PBS containing 10 U/ml heparin at a constant flow rate (∼6 mL/min) until complete clearance of blood was achieved, as indicated by pallor of peripheral organs and clear perfusate. The extracted brains were fixed in 4% PFA for 72 hours, followed by extensive washing and storage in PBS with 0.05% NaN_3_ at 4°C.

### Virus growth kinetics

To assess viral replication kinetics, multi-step growth curves were generated. Briefly, 1 × 10⁶ Na42/13 or BSR-T7/5 cells were infected in suspension at a multiplicity of infection (MOI) of 0.01 in a total volume of 8 ml. Infection was carried out on a roller mixer for 2 h at room temperature. Subsequently, cells were washed three times by centrifugation at 1,500 × g for 5 min and resuspension in 8 ml of the respective cell culture medium. After the final wash, cells were pelleted again at 1,500 × g for 5 min. A 0.5 ml aliquot of the cell culture supernatant was collected as the 0 hours post-infection (hpi) sample and stored at −80 °C. The cell pellet was resuspended, and 100 µl of the cell suspension were seeded together with 100 µl of fresh cell culture medium into a 96-well cell culture plate as an infection control. At 24 hpi, the control plate was fixed and immunostained for RABV nucleoprotein. The remaining cell suspension was transferred to T25 cell culture flasks, adjusted to a final volume of 8.5 ml with cell culture medium, and incubated at 37 °C and 5% CO₂. At 16, 24, 48, 72, and 96 hpi, 0.5 ml supernatant samples were collected, clarified by centrifugation at 1,500 × g for 5 min, and stored at −80 °C. After each sampling, the volume in the T25 flasks was restored to 8.5 ml with fresh cell culture medium. Viral titers were determined by virus titration.

### IFN luciferase assay

A dual-luciferase reporter assay was used to assess effects on promoter activity. Firefly luciferase (FL; *Photinus pyralis*) expression was driven by the promoter of interest, whereas Renilla luciferase (RL; *Renilla reniformis*) was expressed from a constitutive promoter and served as an internal normalization control. Cells were seeded according to the respective experimental setup and co-transfected with FL and RL reporter plasmids at a 100:1 ratio using Polyethylenimine (PEI). The cells in 6-well plates received 1.5 µg FL plasmid and 15 ng RL plasmid per well. For plasmid-transfected samples, cells were washed once with cell culture-grade PBS and lysed after 22 h. Luciferase activities were measured using the Dual-Luciferase® Reporter Assay System according to the manufacturer’s instructions. The samples were analyzed on a Sirius single-sample luminometer, whereas virus-infected samples were measured using an LB 942 TriStar2 plate-reading luminometer. FL activity was recorded after addition of Luciferase Assay Reagent II, followed by quenching and simultaneous initiation of the RL reaction with Stop & Glo® Reagent. For both reactions, luminescence was measured for 10 s after a 2 s delay. FL signals were normalized to RL signals. For analysis of IFN-α-induced ISRE promoter activity in virus-infected cells, Na42/13 cells were first infected with the indicated viruses (MOI = 0.01). At 48 h post-infection, cells were co-transfected with pISRE-Fluc and pCMV-Rluc reporter plasmids, followed by stimulation with 200 U/ml IFN-α for 22–24 h. Luciferase activities were subsequently measured as described above. Statistical significance was assessed by one-way ANOVA followed by Tukey’s multiple-comparisons test.

### Mouse brain tissue extracts and peptide collection for Mass spectrometry

Fixed brain sections juxtaposed to those used for immunofluorescence were placed in 50 µl NuPAGE LDS (Novex by Life Technologies) with 10 mM f.c. DTT (Sigma) and lysed in a tissue lyser (Qiagen) with a 3 mm steel ball at a frequency of 30 Hz for 3 min. The samples were then heated at 100°C for 30 min. The concentration of the samples was quantified by doing a BCA assay (Thermo Scientific), following the manufacturer’s instructions.

For SP3, 25 µg of protein and reconstitution buffer (50 mM HEPES-NaOH pH 8, 100 mM NaCl, 1 mM EDTA pH 8, 0.5% SDC, 1% Triton X-100) was added to a 96-well 200 ml PCR plate (Biozym) at a final volume of 100 µl. Proteins were reduced by adding 10 mM f.c. DTT (Sigma) and heating the plate at 60°C for 30 min shaking at 800 rpm. Proteins were alkylated by adding 25 mM f.c. IAA (Sigma) and incubating in the dark at 25°C for 25 min shaking at 800 rpm. To quench the remaining IAA, 10 mM f.c. DTT was added to the samples followed by incubation at 25°C for 5 min shaking at 800 rpm in a Thermomixer (Eppendorf).

SP3 beads, Sera-Mag Carboxylate-Modified Magnetic Particles (GE Healthcare), were prepared by combining hydrophobic and hydrophilic beads in a 1:1 ratio. The beads were washed two times with 200 µl LCMS-grade H_2_O (VWR). After washing, the beads were resuspended in LCMS-grade H_2_O at a concentration of 50 µg/µl, and 5 µl of beads were added to each sample and mixed by pipetting.

A volume of 150 µl of absolute ethanol was added to the samples + beads and mixed by shaking at 1,000 rpm at 25°C for 15 min. After mixing, the supernatant was removed and the beads were washed three times with 180 µl of 80% LCMS-grade ethanol (Millipore Sigma). After washing, 100 µl of 50 mM ammonium bicarbonate (Sigma) containing 500 ng trypsin (Serva) was added to the samples and incubated overnight at 37°C and 1,000 rpm shaking overnight to digest the proteins.

To collect the peptides, the plate was placed on a magnet (Permagen) for 2 min until the supernatant was clear, and the supernatants were transferred to fresh tubes. To the beads, 100 µl of 0.5% formic acid was added and mixed shaking at 1,000 rpm and 25°C for 10 min. After incubation, the plate was placed on the magnet, and the supernatants were combined with the supernatants previously transferred to fresh tubes in the steps above.

Combined supernatants were desalted via stage-tipping. For StageTips, two layers of Empore C18 (3M) material were pushed into a 200 µl pipette tip. The tips were activated with 50 µl methanol. To wash and equilibrate the tips, 50 µl of Buffer B (80% acetonitrile 0.1% formic acid) was added to the StageTips followed by 50 µl of Buffer A (0.1% formic acid). The supernatants from above were added to the StageTips, and the StageTips were washed again with 50 µl Buffer A.

To elute the peptides from the StageTips, 30 µl of Buffer B was added to the StageTips followed by centrifugation at 500 xg for 2 min with a 24-well PCR plate (Thermo Scientific) for collection. The plate containing the samples was placed in a concentrator (Eppendorf) for 15 min to evaporate the acetonitrile until the final volume was 7 µl. Buffer A was then added to make a final volume of 14 µl. The concentration of the samples was measured using a BCA assay and diluted to 200 ng/µl with Buffer A.

### Mass spectrometry measurement

LC-MS/MS was performed with a NanoElute2 coupled to a TimsTOF-HT (Bruker), and MS data were acquired via Compass HyStar and timsControl. Two microliters of sample was injected and separated on an Aurora Series 3 column (25 cm length, 75 µm inner diameter, 1.7 µm C18 (IonOpticks) in one column mode with a flow rate of 300 nL/min over a 40 min gradient containing a mixture of Buffer A (0.1% formic acid in water) and Buffer B (0.1% formic acid in acetonitrile) described as follows: 2 min 2% Buffer B, 28 min 4% Buffer B, 6 min 17% Buffer B, 1 min 32% Buffer B, and 3 min 95% Buffer. The column temperature was maintained at 50°C via a column toaster (Bruker). The separated peptides were ionized with a CaptiveSpray ion source at a capillary voltage of 1600 V, a dry temp of 180°C, and 3.0 l/min dry gas. MS data were acquired in positive ion and DDA-PASEF mode with 10 PASEF MS/MS scans covering a mass range of 100–1700 m/z.

The raw MS data were analyzed via MaxQuant version 2.4.2.0 with the default settings. Under group-specific parameters, TIMS-DDA was selected. Match between runs was inactivated. Proteins were quantified via the LFQ algorithm (without FastLFQ) with a minimum ratio count of 2 unique peptides. Proteins were identified via database matching with the *Mus musculus* UP000000589_10090 reference database (54,857 entries, UniProt) and a custom rabies fasta files for DogA (8 entries), rSADL16 (6 entries), rCVS11 (5 entries), RABV_DogAP_W265G_M287V_K6 (5 entries), rRABV FoxK9 (5 entries) and rSAD_GasGas (6 entries), with a false discovery rate of 0.01. Contaminants, protein groups identified only by site, reverse database hits, and proteins with fewer than two peptides were removed. Missing values were imputed using a fitted beta-distribution at LOQ.

### Mass spectrometry data analysis

To compare virus concentrations from MS data, the dataset was filtered for viral peptides measured across at least 4 different viruses. The mean LFQ intensities for these peptides were z-scored across the different viruses and plotted within the R environment.

Proteomic depth and data quality were assessed by quantifying the number of identified proteins and peptides (total, razor+unique, and unique) per sample following removal of contaminants and reverse database hits. Reproducibility across biological replicates was evaluated by calculating the coefficient of variation of protein identifications within each virus and timepoint. Missing LFQ values were quantified per sample before and after imputation to assess data completeness and the impact of imputation on downstream analyses. To assess potential batch effects, principal component analysis was performed on imputed log₂ LFQ intensity matrices for each timepoint separately. No systematic relationship between principal component structure and acquisition order was observed.

Differential protein abundance was assessed using log₂ fold changes and corresponding p-values calculated during proteomics data processing. Volcano plots display log₂ fold changes versus −log₁₀-transformed p-values for each virus relative to mock controls.

Volcano plots were generated using precomputed log2 fold changes and corresponding −log10-transformed p-values obtained during proteomics data processing. Fold-change signs were oriented such that positive values indicate higher protein abundance in infected samples relative to mock controls.

Principal component analysis (PCA) was performed on log₂-transformed, imputed LFQ intensity values after centering, removal of reverse hits, potential contaminants and scaling of samples. PCA was applied either to all quantified host proteins (global PCA, including mock samples) or to subsets of proteins enriched relative to mock controls (enriched PCA, excluding mock samples). For visualization and downstream analyses, the first two principal components were used. Virus centroids were defined as the mean PC1 and PC2 coordinates across biological replicates. Pairwise Euclidean distances between virus centroids were computed to quantify inter-virus separation in PCA space. Heatmaps were generated from centroid distance matrices for visualization.

### Immunofluorescence microscopy

Brains were collected following transcardiac perfusion and sectioned into 100 µm-thick coronal slices using a VT1200S vibratome (Leica Microsystems, Fig. 1B). Sections corresponding to Allen Institute Mouse Brain Atlas (https://mouse.brain-map.org/) levels 69–73 were selected for subsequent analyses. The sections were washed with 100% methanol at 1 mL/min for at least 10 hours, followed by 1 hour wash with pure Dichloromethane and further incubated for at least 8 hours before 100% methanol at 0.5 mL/min for 5 hours. The sections were then washed with 30% hydrogen peroxide for 15 min at 1 mL/min, further incubated for at least 8 hours. The samples were rehydrated with sterile distilled water at a flow rate of 2 mL/min for 5 hours, with a further incubation for at least 8 hours. Sections were permeabilized in a buffer containing 0.2% Triton X-100, 20% DMSO, 0.3 M glycine, and 0.02% sodium azide in PBS for 12 hours at 37 °C, followed by blocking with 6% donkey serum, 0.2% Triton X-100 and 10% DMSO in PBS for 8 hours at 37 °C and incubated with primary antibodies for 5 h at 37 °C with shaking at 120 rpm in Innova 42R orbital shaker (Eppendorf). The samples were washed with PBS at a flow rate of 1 mL/min for 12 hours at 37 °C. Sections were then incubated with the secondary antibodies for 5 h at 37 °C with shaking at 120 rpm, protected from light, followed by washing as described above. The samples were mounted on microscope slides in Vectashield and imaged using Leica Stellaris 8 confocal microscope (Leica Microsystems) equipped with HC PL APO 20x/0.75 IMM CORR CS2 objective. 3D mosaics of entire brain sections were recorded and merged with LAS X v.4 software.

### Image analysis

Confocal microscopy records were processed and quantified within Arivis Pro 4.3 (Carl Zeiss AG) software platform. The raw Lif image z-stacks were converted to Sis format using Arivis SIS Converter 3.5.1 (Carl Zeiss AG). Analysis pipelines were implemented using Arivis Pro for image segmentation and object detection with a feature set filtering defined for each immunofluorescence marker. The resulting segmented image stacks were rendered as maximum intensity projections and relevant object feature measurements were exported in table format. In order to perform brain region quantification, manually curated masks of anatomical regions were created according to Mouse Brain Atlas (Allen Institute) for each confocal record in Fiji Software (NIH). The masks were transferred to Arivis Pro software and applied to confocal z-stacks for image segmentation and quantification.

### Gene ontology analysis

Gene Ontology (GO) Biological Process enrichment analysis was performed using the g:Profiler Python package against the *Mus musculus* annotation database. Enrichment was assessed using g:Profiler’s multiple-testing corrected p-values (FDR-adjusted p-values), and GO terms with FDR < 0.05 were considered significantly enriched. To reduce representation of broad or low-confidence categories, enriched terms were filtered to retain GO categories with a minimum term size of ≥10 annotated genes and a minimum intersection size of ≥3 proteins. Enrichment analyses were performed independently for each RABV and infection time point (2, 4, and 6 dpi). Selected immune-associated GO terms were visualized as time-resolved heatmaps using Python, where color intensity represents the significance of enrichment −log₁₀(FDR-adjusted p-value).

### Statistical analysis

Statistical analysis was done using GraphPad Prism 10.4.1 software. Six biological replicates were used in mouse infection for each virus and per time point. Ordinary one-way ANOVA was used unless otherwise stated.

### Ethics statement

The experimental work in mice strictly followed institutional and European guidelines on animal welfare and care according to the Federation of European Laboratory Animal Science Associations (FELASA). Animal experiments were evaluated, reviewed, and approved by the Ethics and Animal Welfare Committee of the Federal State of Mecklenburg-Western Pomerania, Landesamt für Landwirtschaft, Lebensmittelsicherheit und Fischerei Mecklenburg-Vorpommern, 18003 Rostock, Germany (FLI-7221.3-1-032/24). Experimental procedures were supervised by the Institute’s Animal Welfare Officer (FLI) on behalf of the Institutional Animal Care and Use Committee (IACUC).

## Data, code, and materials availability

All data needed to evaluate and reproduce the results in the paper are present in the paper and/or in Zenodo repository (10.5281/zenodo.22768052) and the ProteomeXchange Consortium via the PRIDE repository with the dataset identifier PXD083939. Recombinant RABV generated in this study are available upon request.

## Acknowledgements

We thank Ms. Bärbel Hammerschmidt for assistance in animal infections, Ms. Kathrin Müller, Ms. Katrin Giesow and Ms. Jeanette Kliemt for technical assistance, Mr. Rene Siewert, Ms. Bärbel Berger and Ms. Nicole Sinkwitz for assistance with animal care.

## Author contributions

S.F., F.B. T.M., D.S.U. designed research; S.G., R.M., L.M.Z., C.M.F., L.S., L.R., O.B., F.B., D.S.U. performed research; F.B., S.F., D.S.U. contributed new reagents/analytic tools; S.G., R.M., F.B., D.S.U. analyzed data; and D.S.U. wrote the paper.

