## Supplementary material for "Viral genetic variation programs divergent neuroimmune states during rabies infection": Supplemetal information

**Supplemental information**

| | G | P | N | M | L |
|---|---|---|---|---|---|
| **rFox** | 95.61 | 97.80 | 99.56 | 99.01 | 97.33 |
| **rCVS-11** | 93.32 | 92.93 | 99.33 | 95.05 | 97.56 |
| **rDogA IFN** | 100 | 99.33 | 100 | 100 | 100 |
| **rSAD L16** | 93.89 | 94.28 | 98.67 | 91.58 | 97.18 |
| **rSAD GASGAS** | 93.32 | 94.28 | 98.67 | 91.58 | 97.18 |

**Table S1.** RABV sequence identity (%) in each protein compared to rDogA.

| **Name** | **Characteristics** | **Remark** | **Pathogenicity** | **Reference** | **Accession number** |
|---|---|---|---|---|---|
| rDogA | recombinant field clone | Azerbaijan 2002 | high | 27 | LN879480 |
| rFox | recombinant field clone | Germany 1998 | high | 27 | LN879481 |
| rDogA IFN | interferon-antagonist mutant | rDogA derived - altered P protein (P265G/M287V) | moderate | this study | |
| rCVS-11 | recombinant fixed laboratory clone | | moderate | 28 | LT839616 |
| rSAD L16 | recombinant attenuated RABV live vaccine | | low | 16 | |
| rSAD GASGAS | recombinant 3rd generation attenuated RABV live vaccine | rSAD L16 derived - altered G protein (R333E/N194S) | very low | this study | |

**Table S2.** Characteristics of viruses used in this study.

| Primer | Sequence |
|---|---|
| 1 | 5’-TCCCAGTACACATCTTAAGGGGAGT-3’ |
| 2 | 5’-AGATGTGTACTGGGAGGAGTTGCTTTGGCC-3’ |
| 3 | 5’-GATTTTGCTTAGCTTGTCGGCCTCTAC-3’ |
| 4 | 5’-AAGCTAAGCAAAATCGTACAGGATGATTT-3’ |
| 5 | 5’-AGGGATGTCTTGTGACATTTTTACC-3’ |
| 6 | 5’-ACTTGGATCGTTGAAAGGAC-3’ |
| 7 | 5’-CGTTAATTAACGTCCTTTC-3’ |
| 8 | 5’-GGGAGGATCTCATTCCAAGT-3’ |
| 9 | 5’-ACTAGCTAGCTTATAAAGTGCTGGGTCATCTAAGC-3’ |
| 10 | 5’-TGGATCGTTGAAAGGACGTTAATTAACGTTTACAGTCTGGTCTCACCCC-3’ |
| 11 | 5’-ACCCAGCACTTTATAAGCTAGCTAGTTACAGTCTGGTCTCACC-3’ |

**Table S3.** Primers used to generate recombinant viruses.

| **Peptide name** | **Sequence** |
|---|---|
| **G.1** | **TLMEADAHYK** |
| **M.1** | **ACQLWSDMSLQTQR** |

| | |
|---|---|
| **N.1** | **ELQEYEAAELTK** |
| **N.2** | **FLAGTYDMFFSR** |
| **N.3** | **IEQIFETAPFVK** |
| **N.4** | **MDADKIVFK** |
| **N.5** | **MFEPGQETAVPHSYFIHFR** |
| **N.6** | **SPEAVYTR** |
| **P.1** | **IFVNPSAIR** |
| **P.2** | **CVLGWVALANSK** |
| **P.3** | **IMQDDLNR** |

**Table S4.** Amino acid sequences of detected RABV peptides.

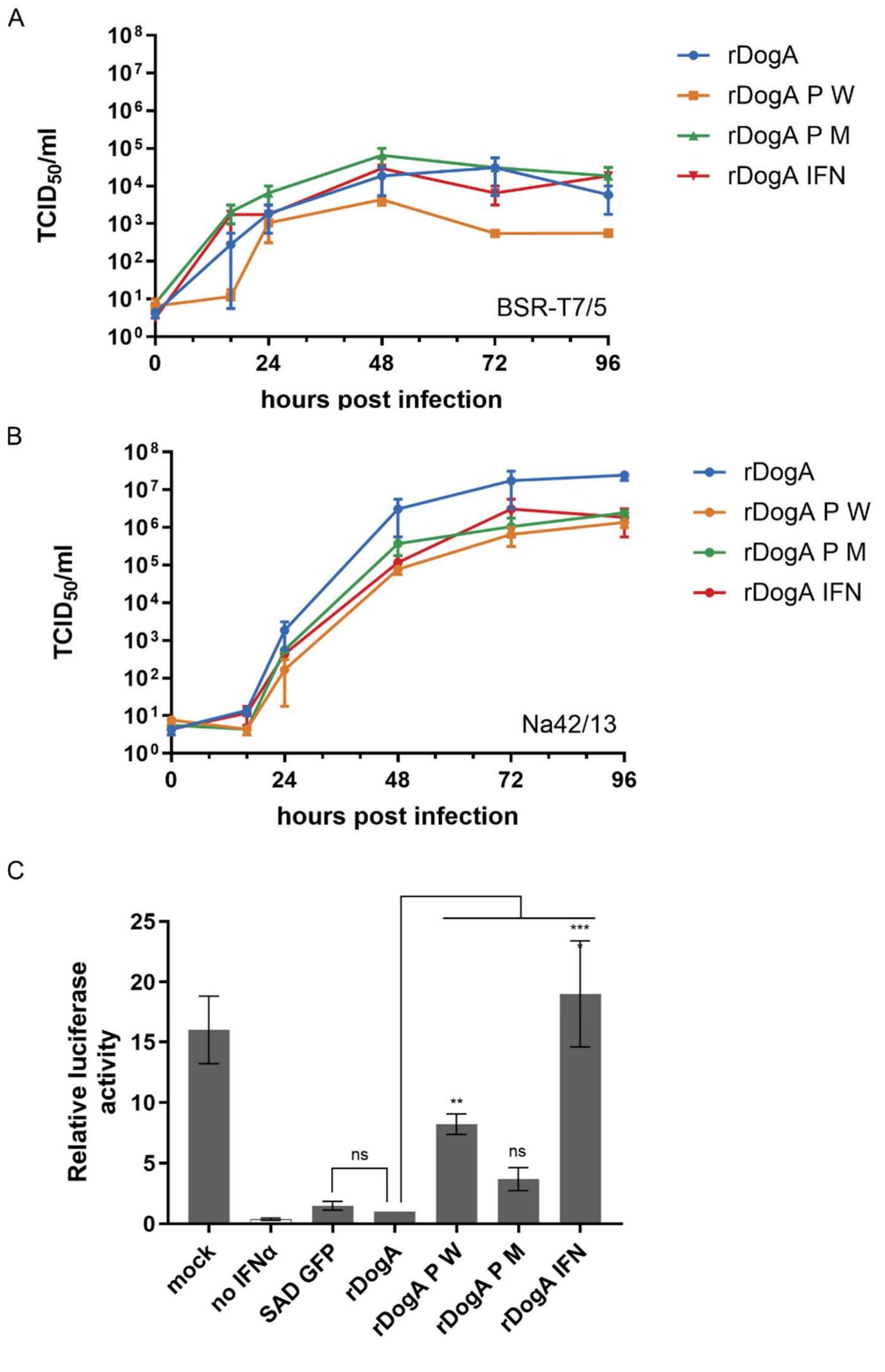


**Figure S1.** Combined introduction of the W265G and M287V mutations in field virus P proteins results in loss of type-I IFN antagonistic activity. (A and B) Replication kinetics of wild type and mutant viruses in Na 42/13 and BsrT7/5 cells. Shown is the mean of two independent experiments. (C) Na42/13 cells were infected (MOI = 0.01) with the indicated viruses, incubated for 48 h followed by co-transfection of pISRE-Fluc and pCMV-Rluc, and induction with 200 U/ml IFN-α for 22-24 h. Mean ± SD of three independent experiments are shown. ns: not significant; ****p ≤ 0.0001.

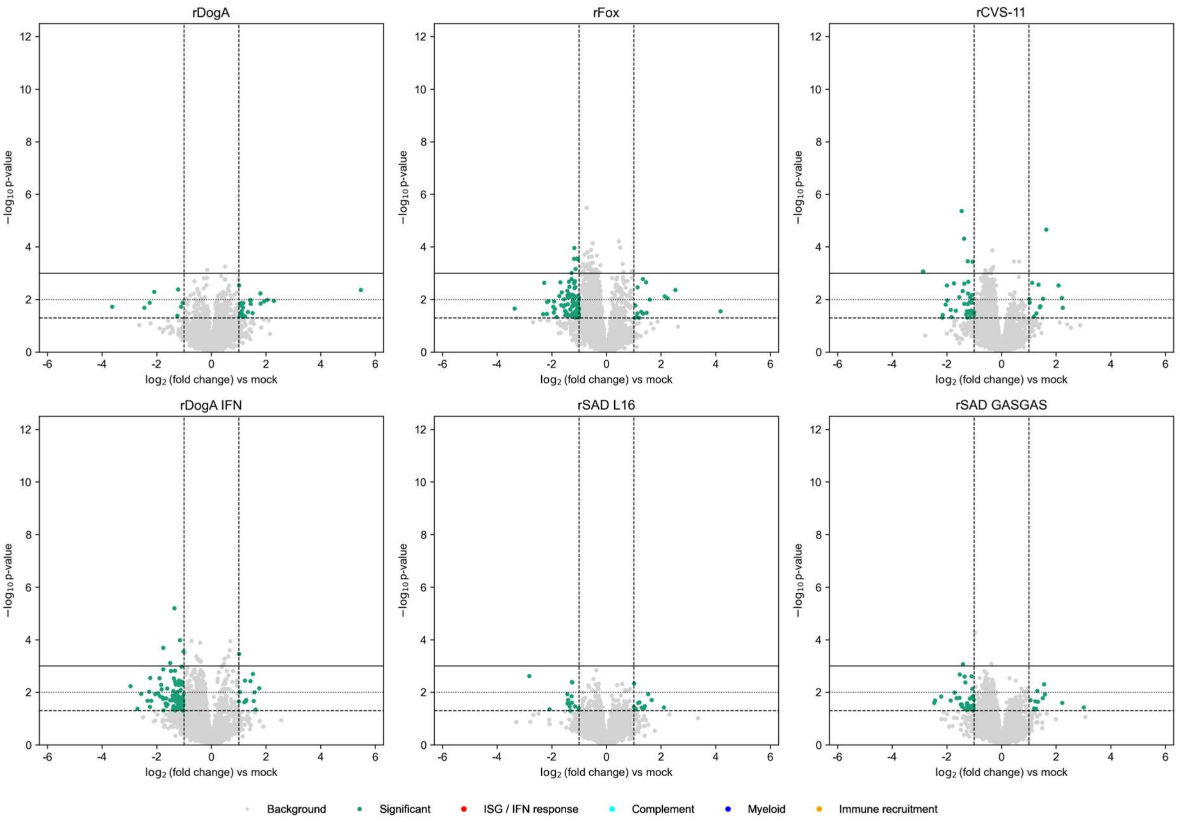


**Figure S2.** Volcano plots showing protein changes in each virus compared to mock at 2 dpi. Each dot represents a protein, with highlighted significant $\log_2$ fold changes ($\geq 1$) and $-\log_{10}$ p-values ($p < 0.05$).

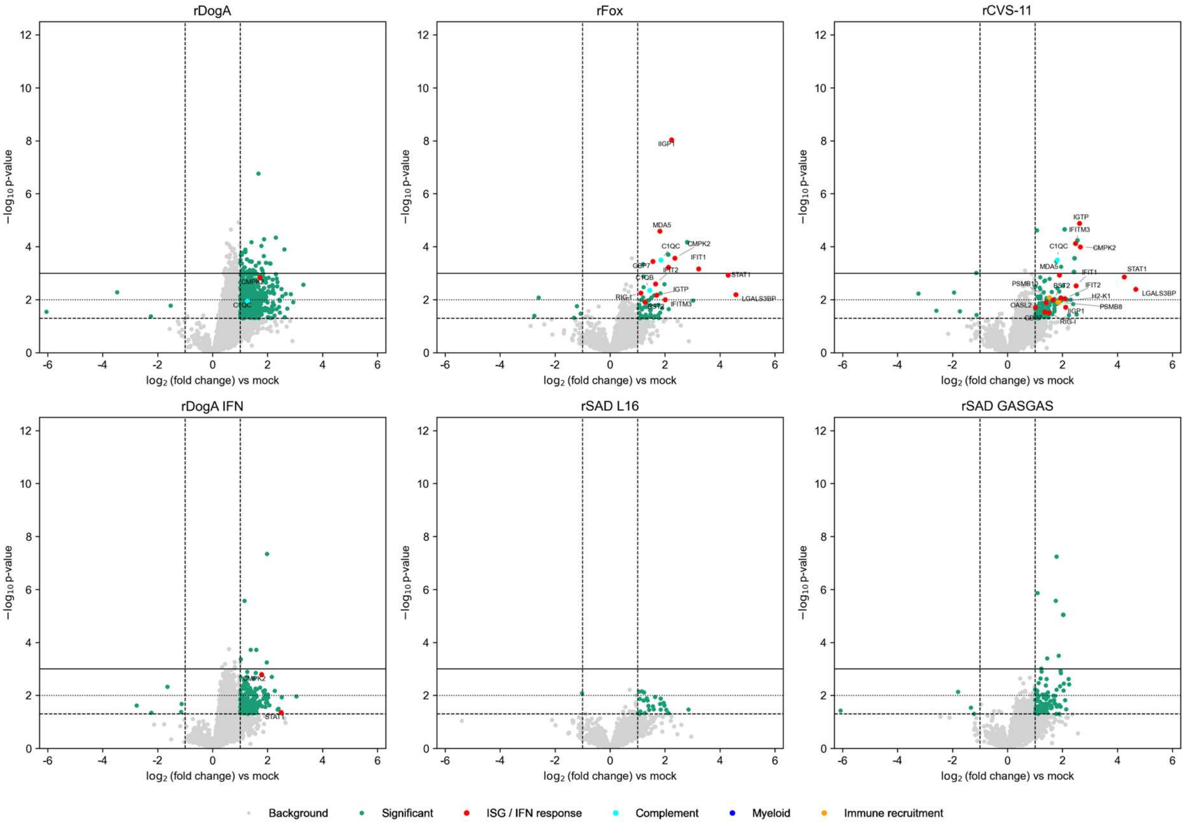

**Figure S3.** Volcano plots showing protein changes in each virus compared to mock at 4 dpi. Each dot represents a protein, with highlighted significant $\log_2$ fold changes ($\geq 1$) and $-\log_{10}$ p-values ($p < 0.05$).

| Peptide name | Peptide sequence |
|---|---|
| G.1 | TLMEADAHYK |
| M.1 | ACQLWSDMSLQTQR |
| N.1 | ELQEYEAAELTK |
| N.2 | FLAGTYDMFFSR |
| N.3 | IEQIFETAPFVK |
| N.4 | MDADKIVFK |
| N.5 | MFEPGQETAVPHSYFIHFR |
| N.6 | SPEAVYTR |
| P.1 | IFVNPSAIR |
| P.2 | CVLGWVALANSK |
| P.3 | IMQDDLNR |

**Table S4.** Amino acid sequences of RABV peptides detected by mass spectrometry.

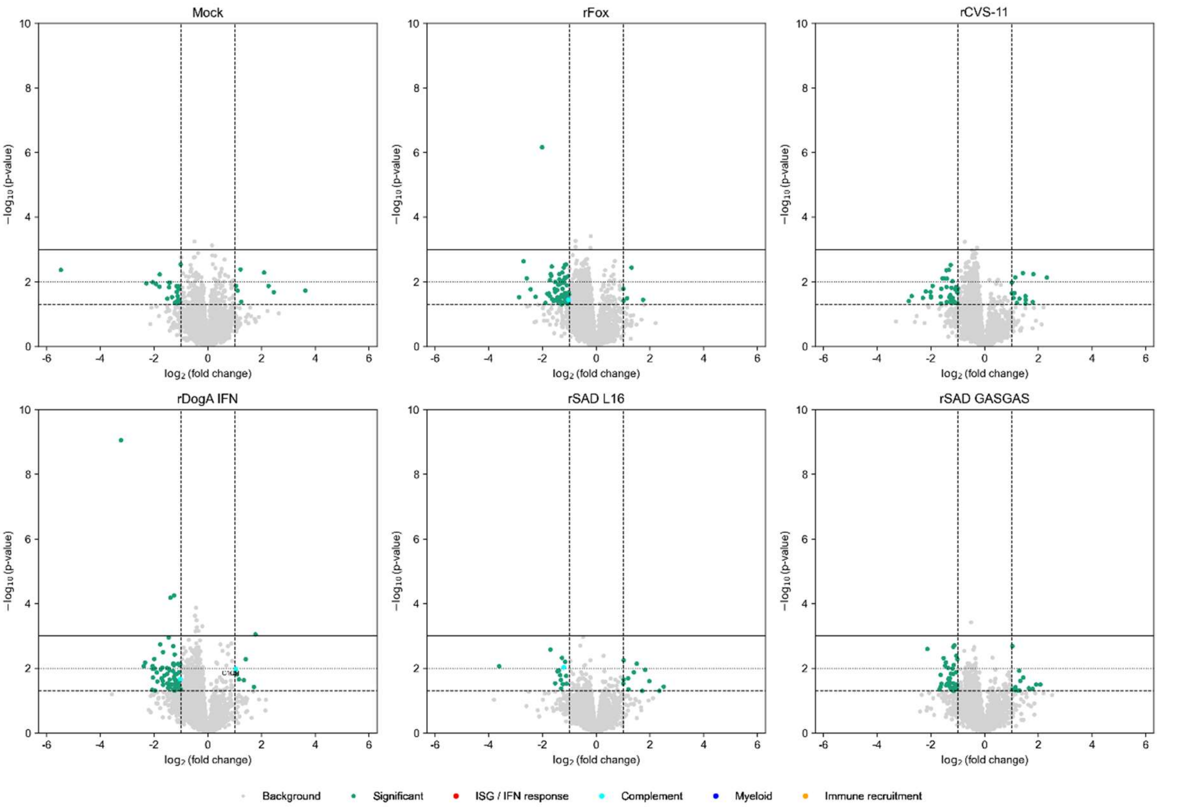


**Figure S4.** Volcano plots showing protein changes in each virus compared to rDogA at 2 dpi. Each dot represents a protein, with highlighted significant $\log_2$ fold changes (≥ 1) and $-\log_{10}$ p-values ($p < 0.05$).

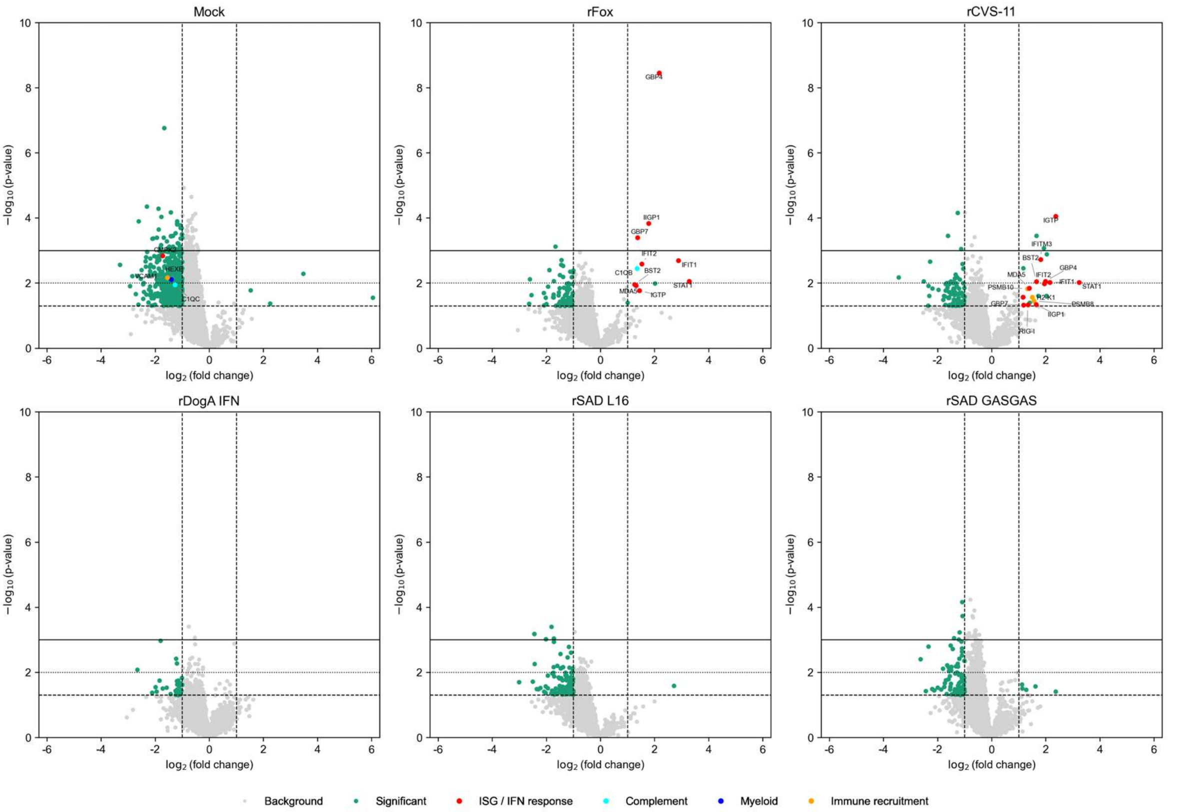

**Figure S5.** Volcano plots showing protein changes in each virus compared to rDogA at 4 dpi. Each dot represents a protein, with highlighted significant $log_2$ fold changes (≥ 1) and $–log_{10}$ p-values ($p < 0.05$).

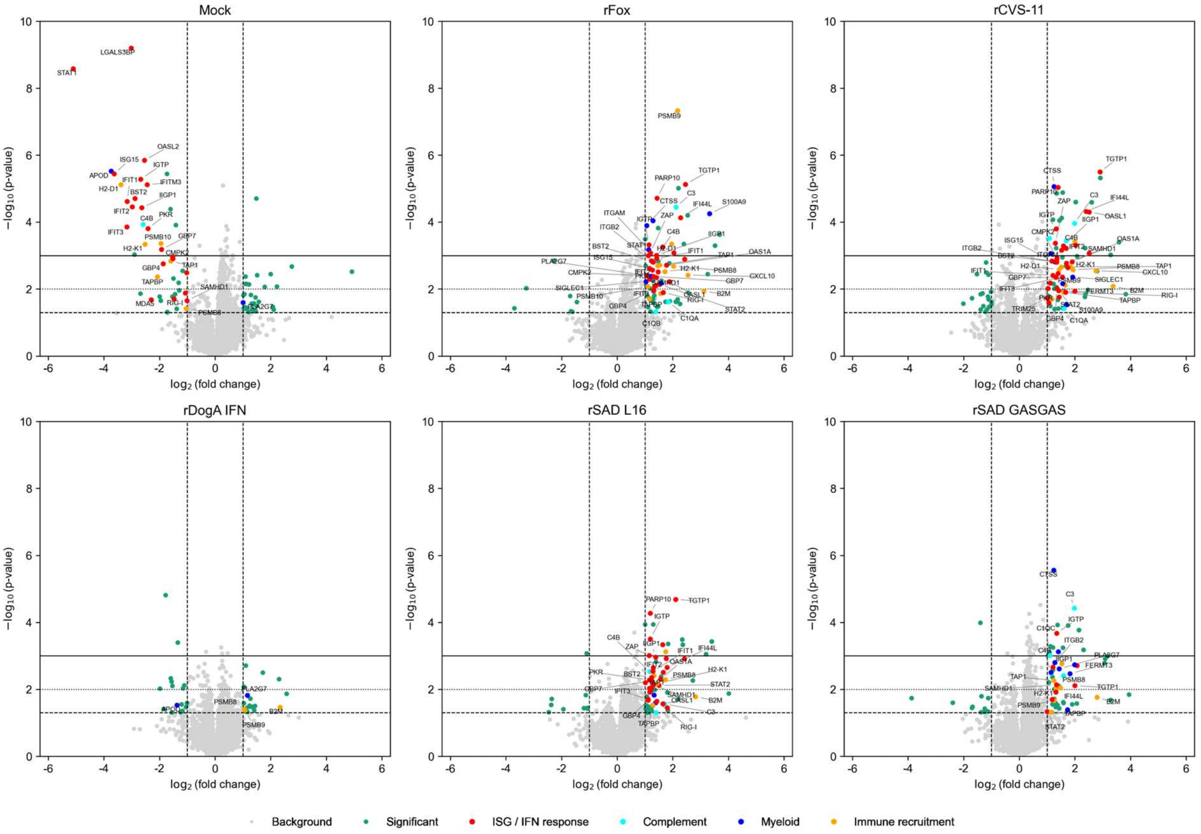


**Figure S6.** Volcano plots showing protein changes in each virus compared to rDogA at 6 dpi. Each dot represents a protein, with highlighted significant $log_2$ fold changes (≥ 1) and $–log_{10}$ p-values ($p < 0.05$).

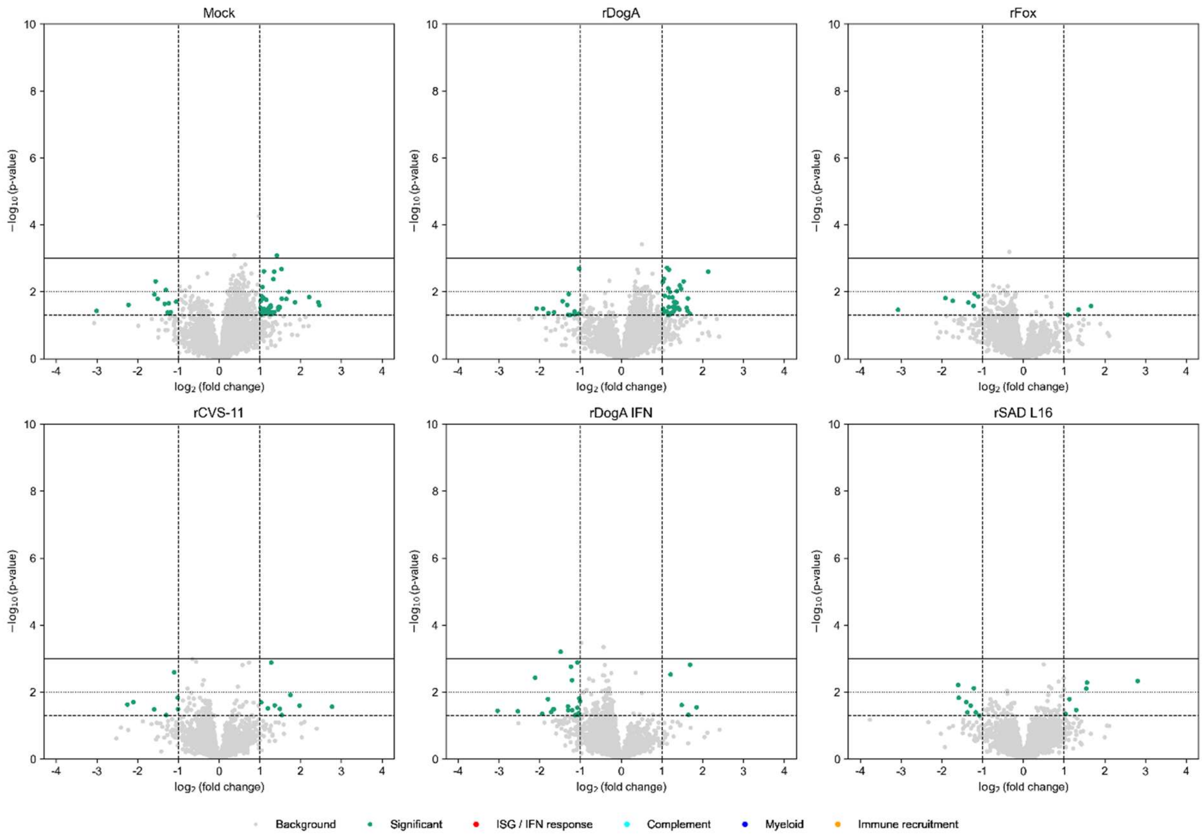


**Figure S7.** Volcano plots showing protein changes in each virus compared to rSAD GASGAS at 2 dpi. Each dot represents a protein, with highlighted significant $\log_2$ fold changes (≥ 1) and $-\log_{10}$ p-values ($p < 0.05$).

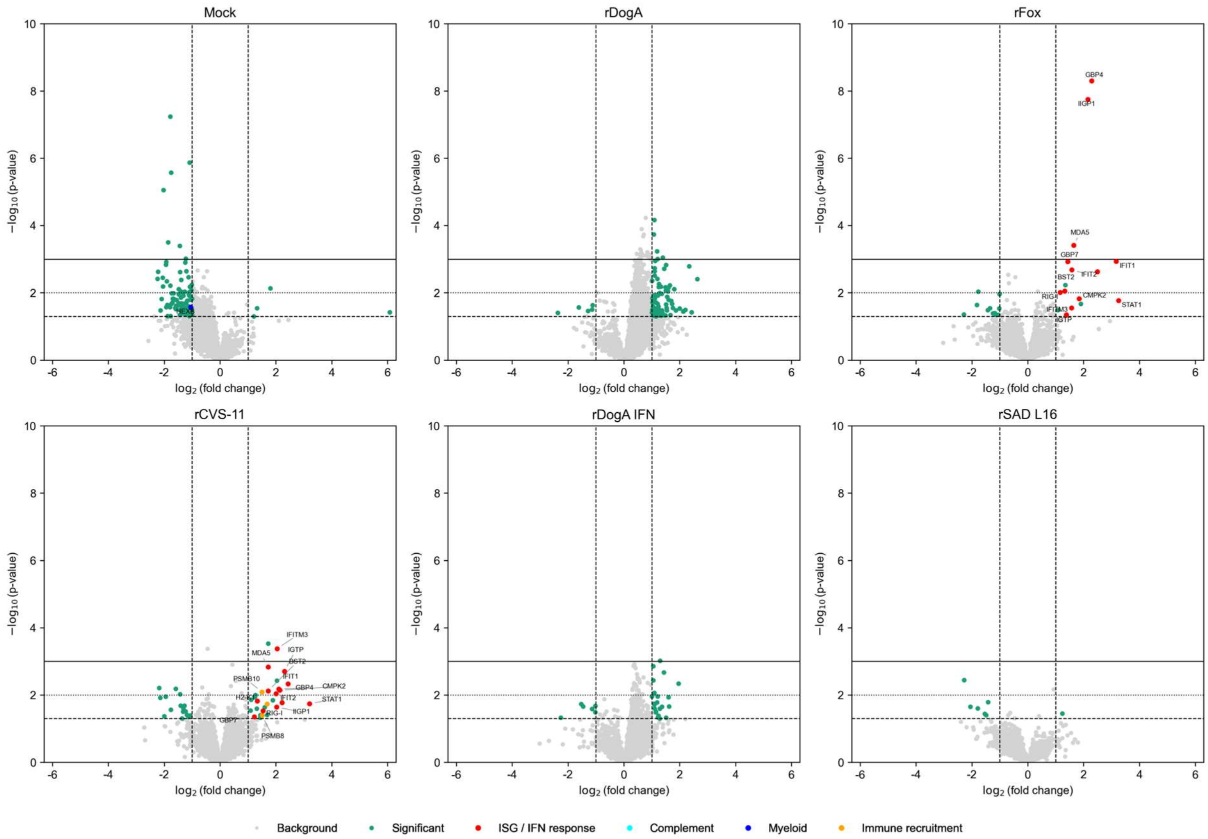

**Figure S8.** Volcano plots showing protein changes in each virus compared to rSAD GASGAS at 4 dpi. Each dot represents a protein, with highlighted significant $\log_2$ fold changes (≥ 1) and $-\log_{10}$ p-values ($p < 0.05$).

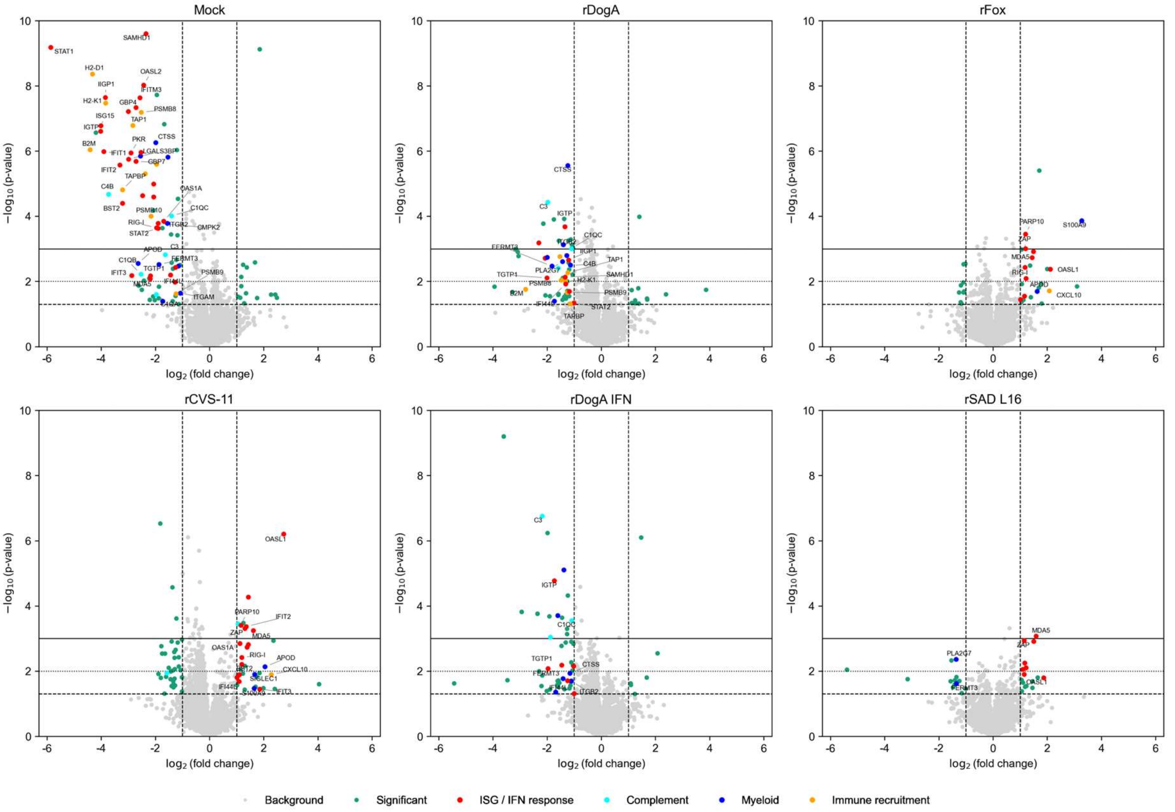


**Figure S9.** Volcano plots showing protein changes in each virus compared to rSAD GASGAS at 6 dpi. Each dot represents a protein, with highlighted significant $\log_2$ fold changes (≥ 1) and $-\log_{10}$ p-values ($p < 0.05$).

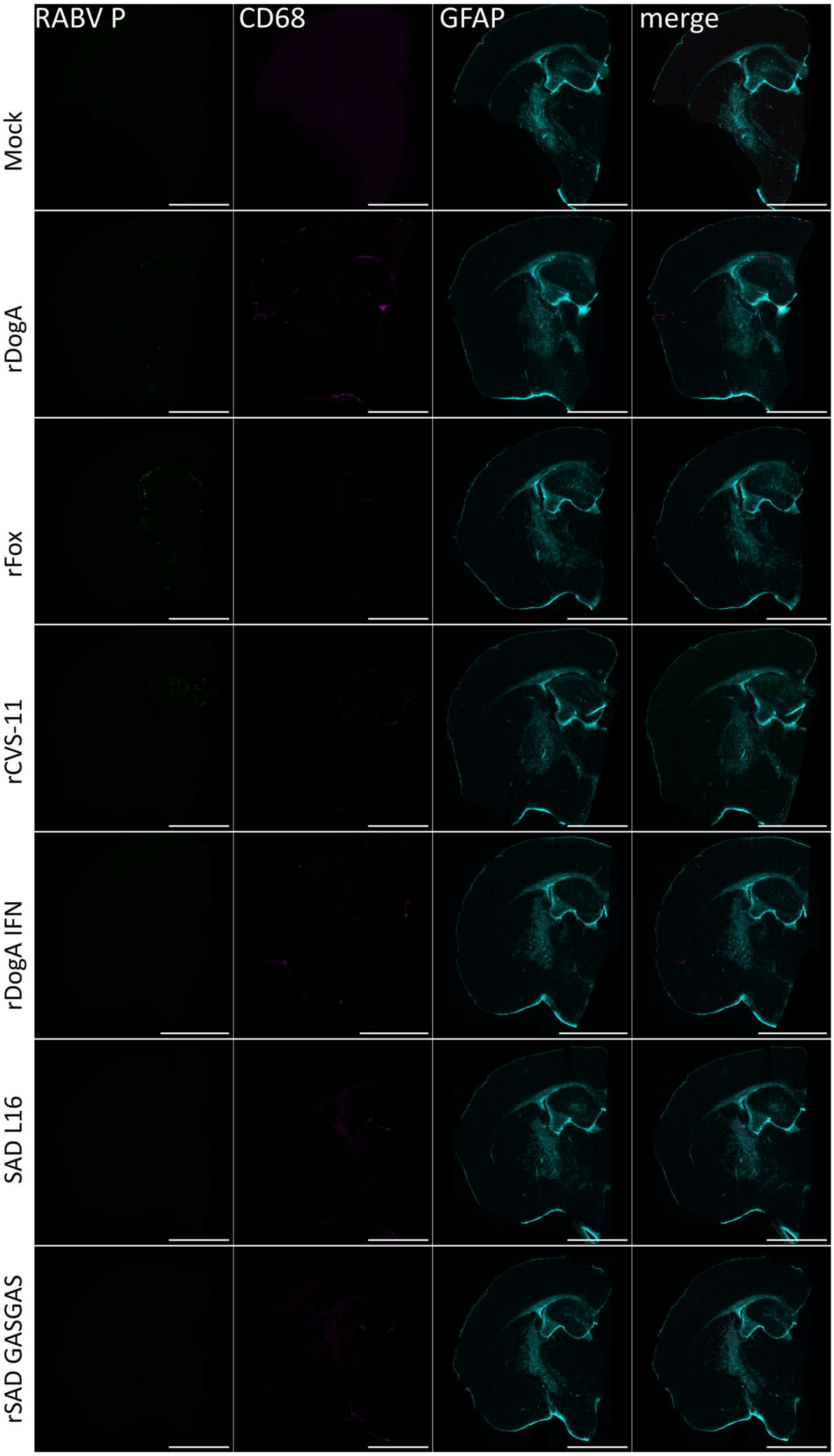


**Figure S10.** Spatial profiling of infection and neuroimmune response in mouse brain at 2 dpi. Representative confocal images of brain sections showing RABV P, CD68 and GFAP in each RABV.

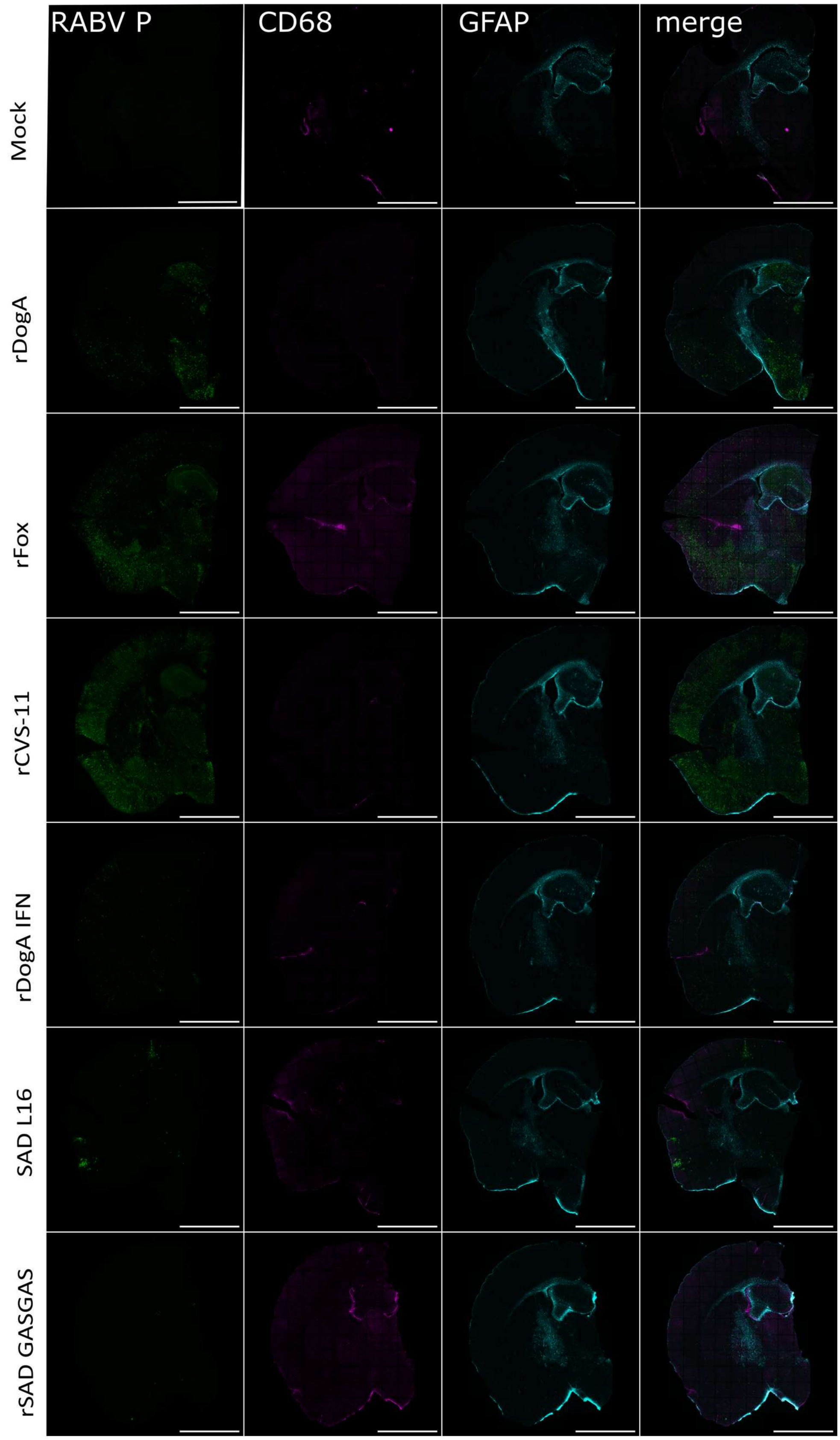


**Figure S11.** Spatial profiling of infection and neuroimmune response in mouse brain at 4 dpi. Representative confocal images of brain sections showing RABV P, CD68 and GFAP in each RABV.

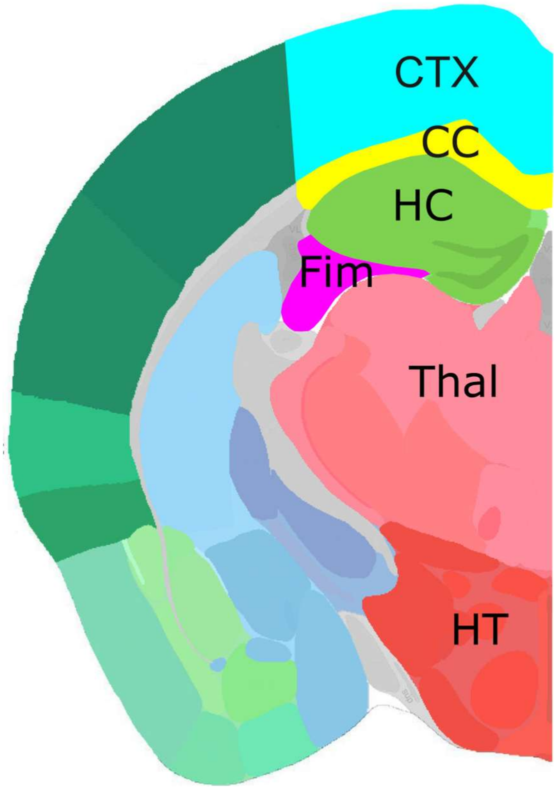


**Figure S12**. Schematic diagram showing quantified brain regions (modified from Allen mouse brain atlas). CTX – Cerebral cortex, CC – Corpus callosum, HC – Hippocampus, Fim – Fimbria, Thal – Thalamus, HT – Hypotalamus.

### Cerebral cortex

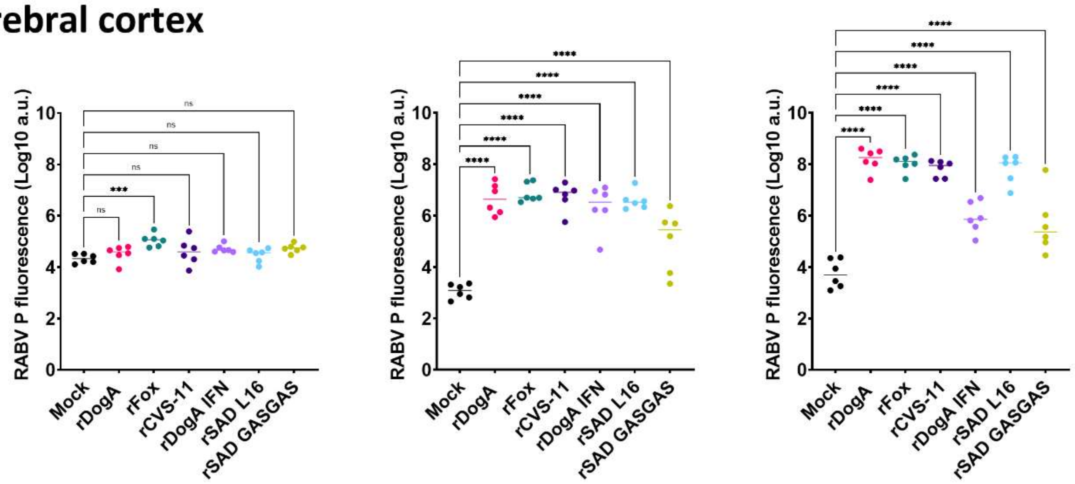


### Corpus callosum

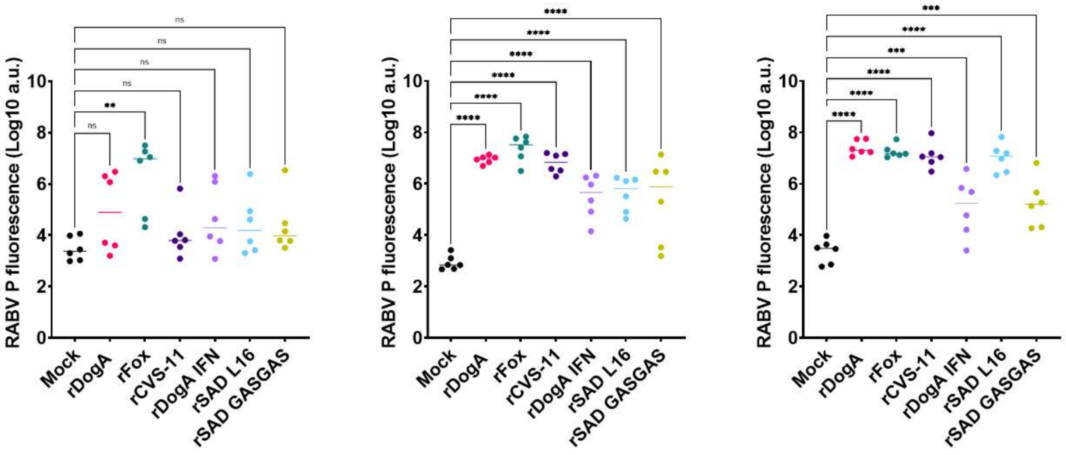


### Hippocampus

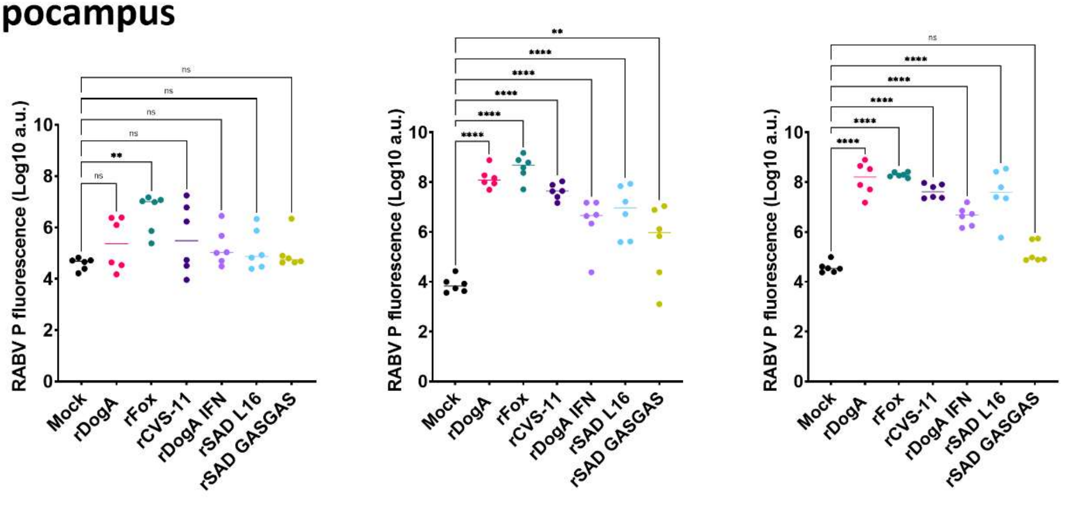


**Figure S13**. Quantitative analysis of RABV P fluorescence levels in Fimbria, Thalamus and Hypothalamus. Points corresponding to individual animals and median are shown. Ordinary one-way ANOVA was used to compare infected groups with mock controls. *P < 0.05, **P < 0.01 ***P < 0.001****P < 0.0001.

### Fimbria

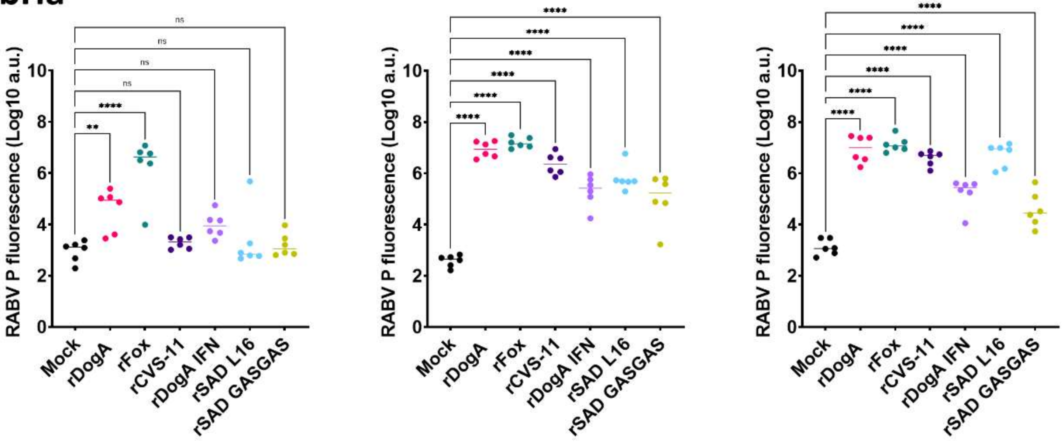


### Thalamus

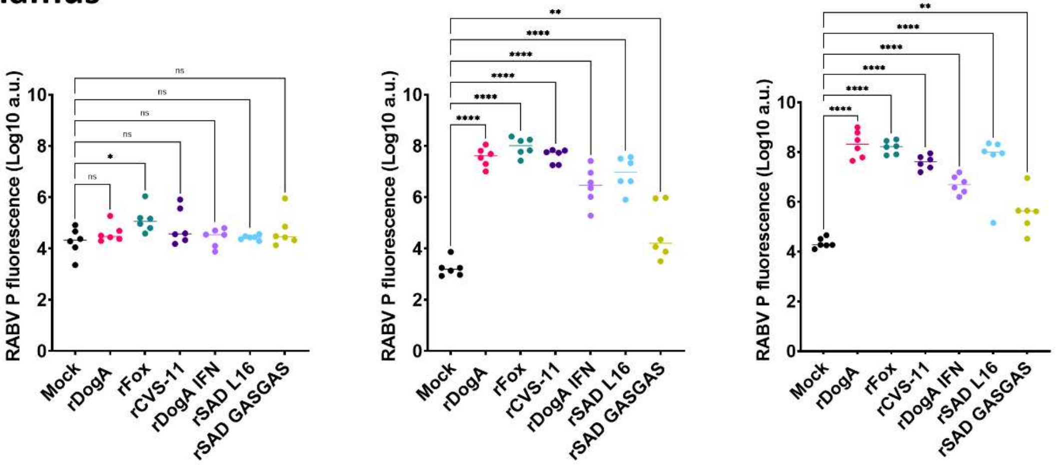


### Hypothalamus

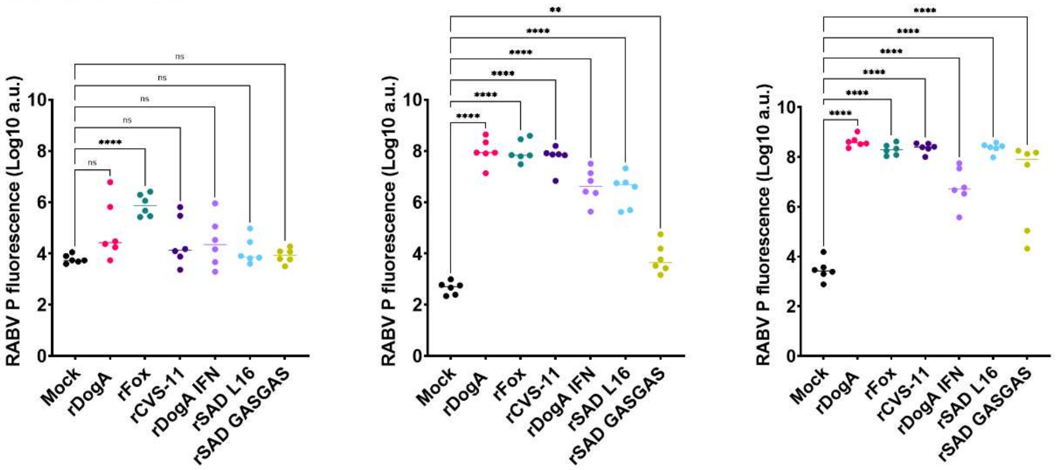


**Figure S14**. Quantitative analysis of RABV P fluorescence levels in Cerebral cortex, Corpus callosum and Hippocampus. Points corresponding to individual animals and median are shown. Ordinary one-way ANOVA was used to compare infected groups with mock controls. *P < 0.05, **P < 0.01 ***P < 0.001****P < 0.0001.

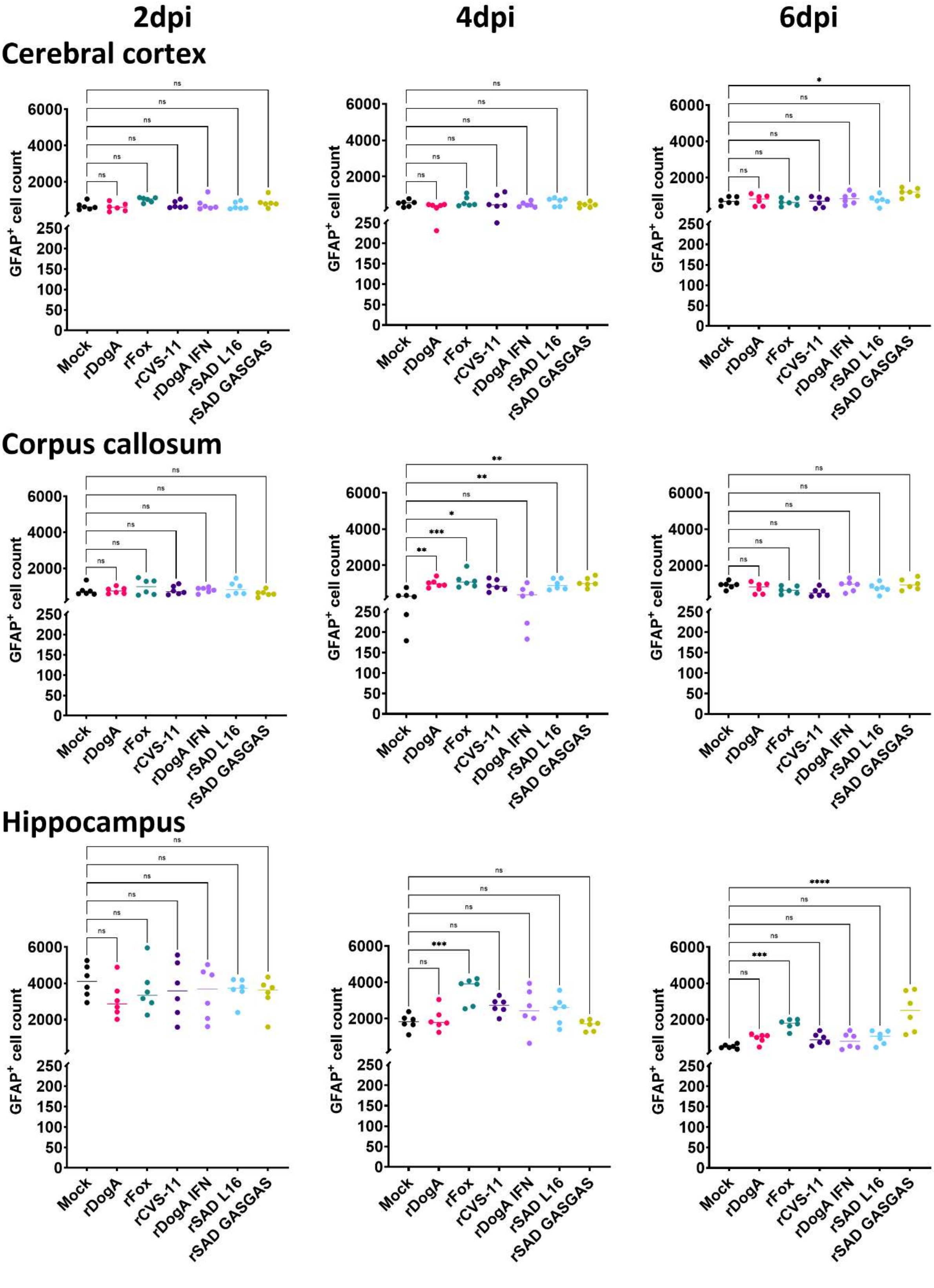


**Figure S15**. Quantitative analysis of GFAP fluorescence levels in Fimbria, Thalamus and Hypothalamus. Points corresponding to individual animals and median are shown. Ordinary one-way ANOVA was used to compare infected groups with mock controls. *P < 0.05, **P < 0.01 ***P < 0.001****P < 0.0001.

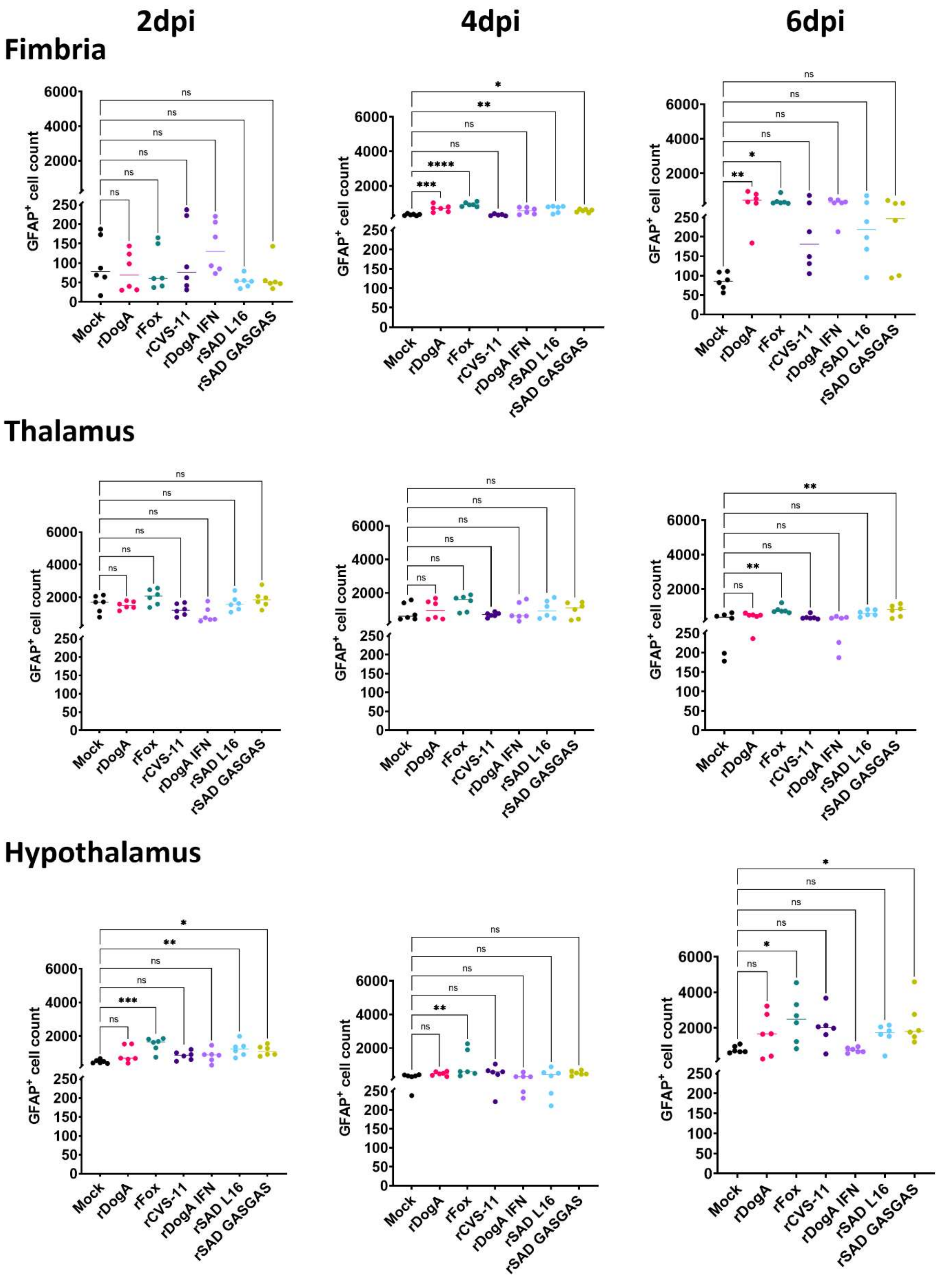


**Figure S16**. Quantitative analysis of GFAP fluorescence levels in Cerebral cortex, Corpus callosum and Hippocampus. Points corresponding to individual animals and median are shown. Ordinary one-way ANOVA was used to compare infected groups with mock controls. *P < 0.05, **P < 0.01 ***P < 0.001****P < 0.0001.

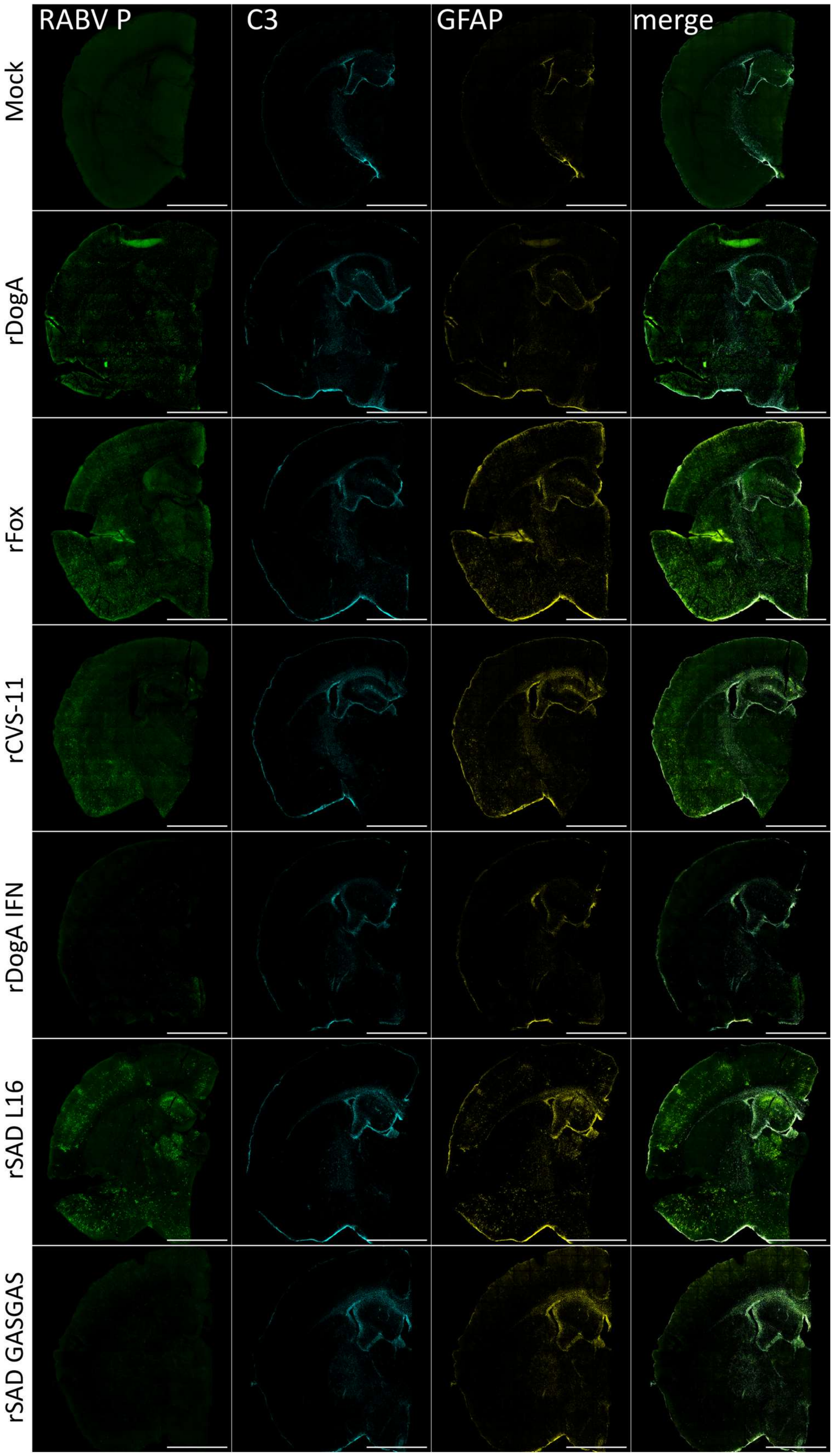


**Figure S17.** Spatial profiling of C3 at 6 dpi. Representative confocal images of brain sections at 6 dpi showing RABV P, C3 and GFAP in each RABV.

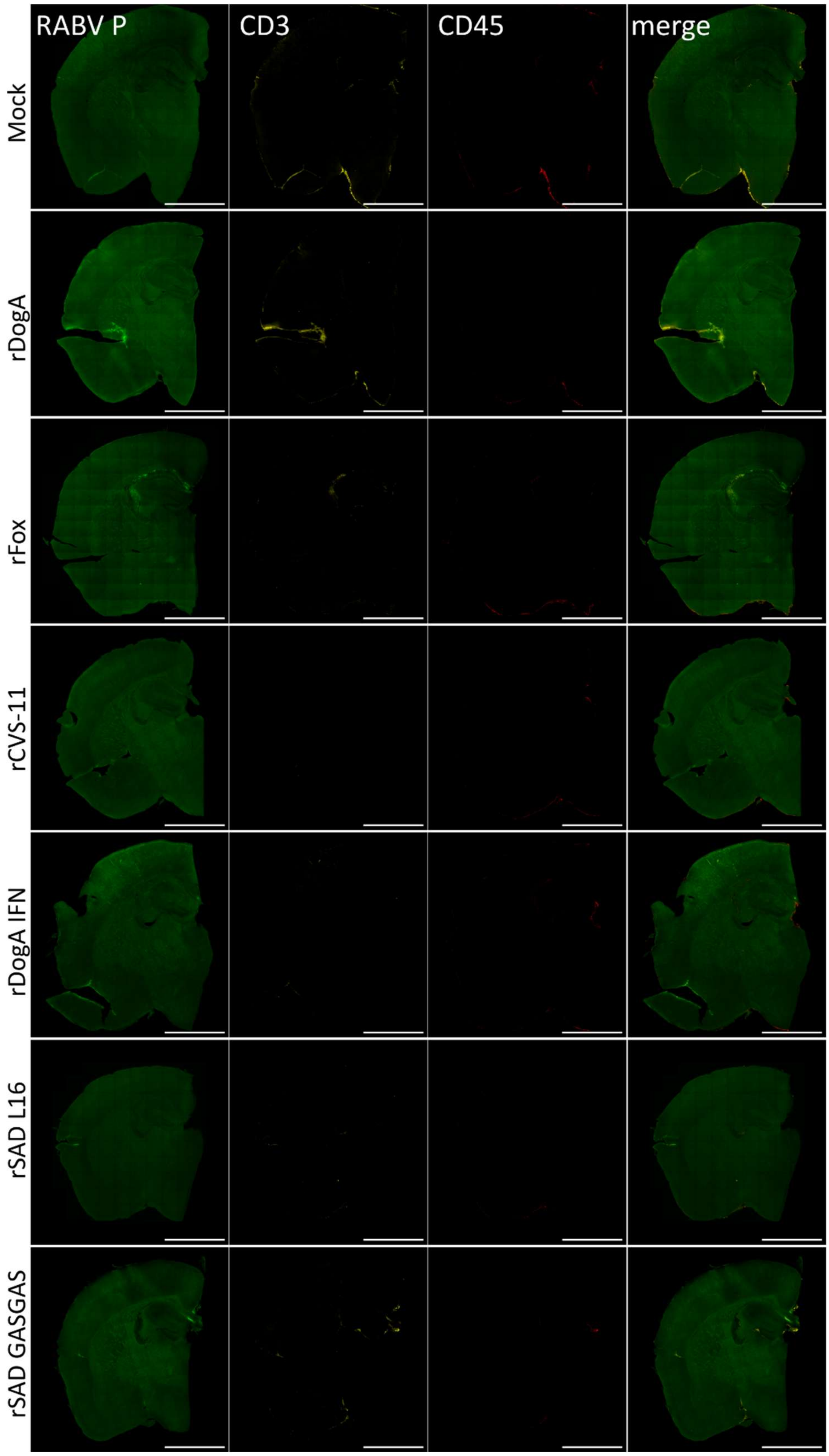


**Figure S18.** Spatial profiling of T cell response following RABV infection at 2 dpi. Representative confocal images of brain sections showing RABV P, CD3 and CD45 in each RABV.

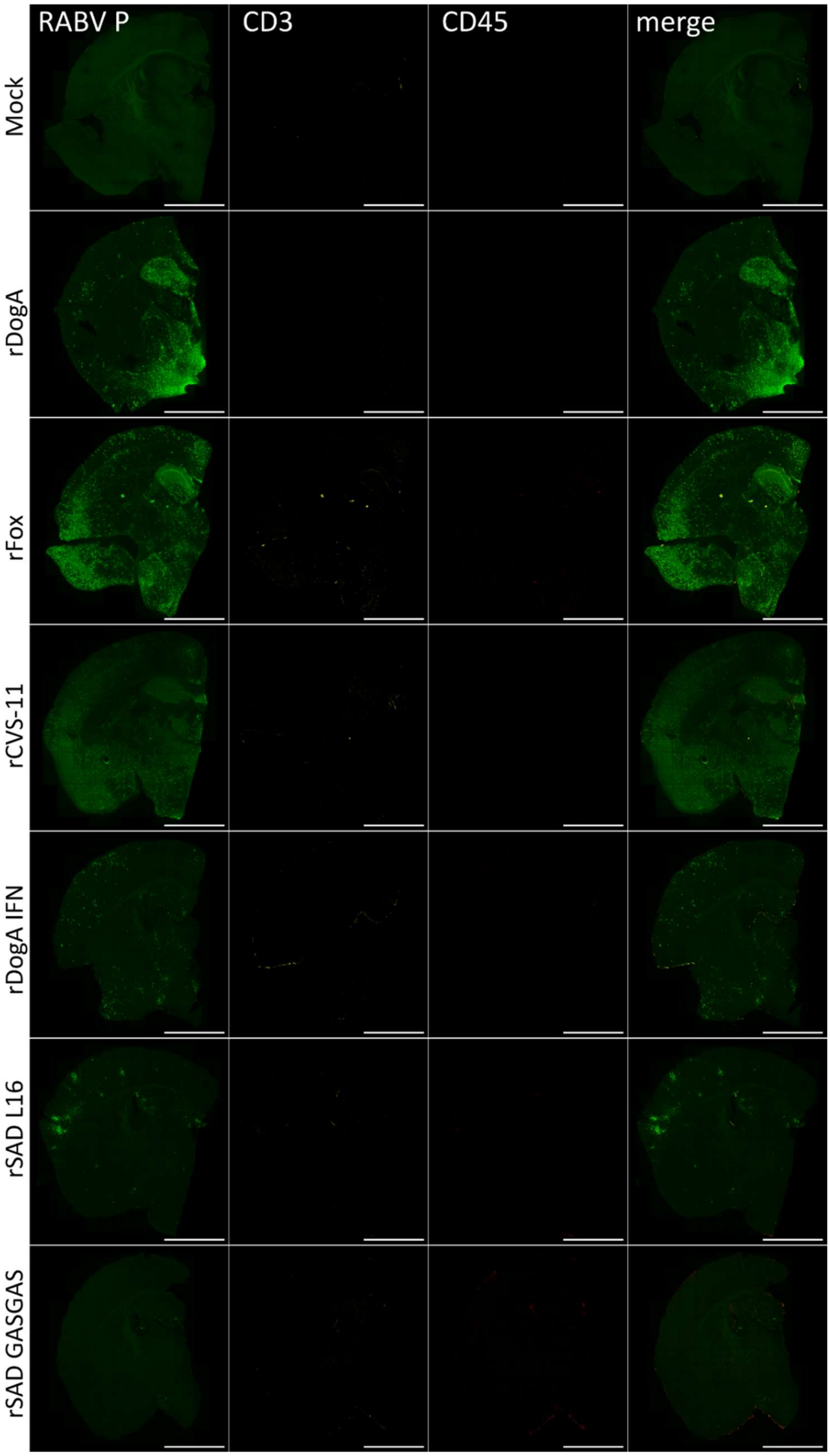


**Figure S19.** Spatial profiling of T cell response following RABV infection at 4 dpi. Representative confocal images of brain sections showing RABV P, CD3 and CD45 in each RABV.

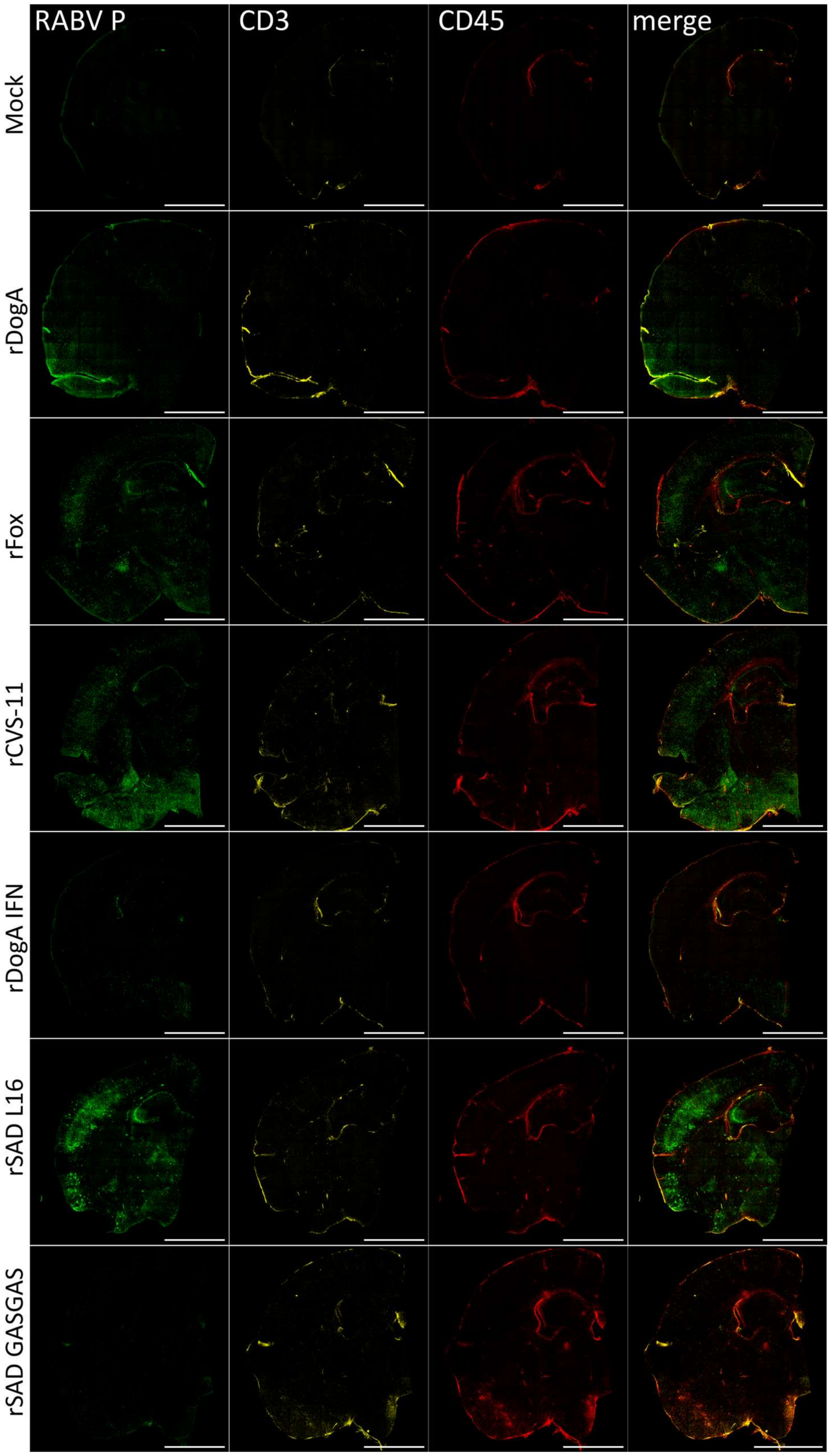


**Figure S20.** Spatial profiling of T cell response following RABV infection at 6 dpi. Representative confocal images of brain sections showing RABV P, CD3 and CD45 in each RABV.
